# Repeated Evolution of Sociality is Associated with Reduced Mutation Rates in Spiders

**DOI:** 10.1101/2025.07.09.663850

**Authors:** Jilong Ma, Anne Aagaard, Jesper Bechsgaard, Anja Ebert, Peter Sørud Porsborg, Iva Kovacic, Sarah Mak, Joseph Nesme, M. Thomas P Gilbert, Peter Andersen, Palle Villesen, Trine Bilde, Mikkel Heide Schierup

## Abstract

Germline mutation rates influence the pace of molecular evolution, yet the roles of selection and life history in shaping their evolution remain to be determined. Comparative systems with replicated evolutionary transitions provide a unique opportunity to determine how changes in life history influence germline mutation rates. In the spider genus *Stegodyphus,* permanent sociality evolved independently three times within the past million years and is associated with obligate inbreeding, female-biased sex ratios, reduced fecundity, and sharply reduced effective population sizes. We sequenced 202 parent-offspring trios from 34 full-sibling families across three social and four closely related subsocial species and analysed quality-filtered trios in a phylogenetic comparative framework. Each independent transition to sociality was associated with an approximately 2-fold reduction in the *de novo* mutation rate in the germline. Phylogenetic analyses of synonymous branch lengths suggest that the mutation rates declined in parallel with the transitions to sociality. These rapid reductions in mutation rates in social lineages with small effective population sizes run counter to the drift-barrier hypothesis, which predicts that reduced selection efficacy would lead to higher mutation rates. We find no evidence that reduced mutation rates in the social species was favoured by selection for improving DNA repair efficiency, since there is no upregulation of DNA repair pathway genes in the ovaries of the social species. On the contrary, the mutation rate is reduced across mutational classes and in somatic tissue in social species compared with their subsocial counterparts. These patterns suggest that the reduction in mutation rate in social spiders is a consequence of convergent life history changes, including reduced body size and production of fewer, larger eggs. Our results highlight that the evolution of sociality, which entails major life history changes, can rapidly reshape fundamental evolutionary parameters, such as the germline mutation rate.

## Main

The mutational process refers to the biological mechanisms and dynamics by which mutations arise, accumulate, and vary across genomes and over time (*1–7*). It is the primary source of genetic diversity, which fuels the evolution of functional traits by enabling both adaptation and the accumulation of genetic load, ultimately affecting the long-term persistence of populations. Studies in vertebrates have shown that the mutational process is broadly conserved across species (*5*), with mutation rates per generation typically varying by no more than one order of magnitude in mammals (*4*, *8–13*), fish (*14*), reptiles (*5*), and birds (*5*). Variation in life history, such as generation time, accounts for a significant portion of the difference in mutation rate among species (*5*, *15*, *16*). Furthermore, species with larger effective population sizes (*N_e_*) tend to have lower mutation rates per generation, as natural selection is more effective at maintaining accurate and efficient DNA repair systems in larger populations, and mutations are more likely to be deleterious than advantageous (*17*, *18*). This pattern is commonly explained by the drift-barrier hypothesis (*19*), which proposes that the extent to which selection can reduce mutation rates is constrained by the opposing force of genetic drift. However, life history traits and *N_e_* often covary across large phylogenetic scales: large, long-lived animals with longer generation time tend to have smaller *N_e_* than small, short-lived organisms. This covariance has made it difficult to determine the relative impact of life history traits and *N_e_* (*15*, *16*). Finally, environmental factors such as UV exposure and heat can influence mutation rates, depending on how effectively cells are protected from external damage, as demonstrated by studies of somatic mutations in tissues like the skin, respiratory mucosa, and digestive tract (*20–23*). In primates, around 70% of the variation in germline mutation rate among individuals is explained by parental age, particularly paternal age (*13*, *24*, *25*), indicating a minimal effect of the environment (*26*). Mammals generally exhibit relatively stable mutation rates across species compared to birds and reptiles (*5*), probably reflecting conserved biological constraints and shared life history traits.

Comparatively less is known about germline mutation rates in invertebrates. Estimates, mostly from *Drosophila* and a few other insects, show that the germline mutation rate per generation (arthropod mean of 2.82e-09) is lower than that of vertebrates (mammals mean of 8.36e-09) (*27*). Invertebrates differ from vertebrates in traits that may shape how mutation rates evolve and vary. Invertebrates occupy an exceptionally broad life history space including body size, reproductive mode, developmental strategy and offspring investment (*28*), which may predict greater variation in mutation rates among taxa, if life history traits are major drivers of mutation rate variation as has recently been proposed (*16*). Additionally, their relatively smaller body sizes and more exposed germline or early development stages may make individual mutation rates more sensitive to environmental stressors. Indeed, thermal stress was shown to increase the mutation rate in *Drosophila* (*29*, *30*), the harlequin fly (*Chironomus riparius*) (*31*, *32*) and silkworms (*Bombyx mori*) (*33*). Thus, invertebrates provide a powerful but still underexplored system for testing how *N_e_* and life history jointly shape germline mutation rate variation.

Here, we present a comparative study of germline mutation rate evolution in spiders from the genus *Stegodyphus*, which exhibit contrasting life history traits and effective population sizes predicted to influence their mutational processes. Across the *Stegodyphus* phylogeny, we find repeated and independent major evolutionary transitions from a solitary and subsocial, outbreeding lifestyle to a highly inbreeding, permanently social lifestyle (*34–37*). Permanent sociality has evolved via the subsocial route (*36*, *38*): subsocial species exhibit extended maternal care, including suicidal maternal sacrifice (matriphagy), in which the mother is consumed by her offspring, followed by communal juvenile living before dispersing prior to mating to live and reproduce solitarily (*35*, *39*). Convergent transitions to sociality happen by the elimination of premating dispersal and a shift to reproduction within family groups (*35*). These repeated social transitions show convergent and highly similar life history changes: 1) Individuals become smaller; 2) the primary sex ratio changes from equal to a strong female bias; 3) transitions from outcrossing to inbreeding mating systems; 4) females lay fewer but larger eggs; and 5) social spiders evolve cooperative breeding where only a fraction of the females reproduce and the remaining females engage in brood care as non-reproducing helpers in the communal nest (*39–45*). These life history transitions cause dramatically lower effective population sizes of the social species by more than 20-fold reductions in *N_e_* after social transitions (*46–48*).

The genus *Stegodyphus* includes three independently and recently derived social lineages, each with a closely related subsocial sister species, offering a unique comparative framework for investigating the genomic consequences of convergent transitions to sociality (*35*, *36*, *47*, *49*). Phylogenetic analyses show that social lineages form short terminal branches, suggesting that sociality may represent an evolutionary dead-end (*36*, *46*). Indeed, comparative genomic analyses reveal evidence of genomic decline through the accumulation of deleterious mutations in social lineages compared to their subsocial sister species (*46*, *47*, *49*, *50*). This raises the question of how mutation rates respond to social transitions: social species could have evolved a lower mutation rate, which reduces the risk of accumulating harmful mutations, either as a consequence of life history changes or as an evolutionary response to the increased selection efficacy against mutator alleles due to the reduced effective recombination rate with high levels of inbreeding (*51*, *52*). Alternatively, social species could experience a higher mutation rate, as predicted by the drift barrier hypothesis, in response to strong reduction of genome-wide selection efficacy due to lower *N_e_*. To address these questions, we performed a large-scale parent-offspring trio sequencing study in seven species of *Stegodyphus* spiders to determine the consequences of three convergent transitions to sociality on the germline mutation process relative to their non-social sister species.

## Results and Discussion

### Consistent reduction in germline mutation rates in social spider species

We estimated the mutation rate in seven *Stegodyphus* species, representing three pairs of social and subsocial sister species, as well as one outgroup subsocial species. We did this by mapping short-read sequences from parent-offspring trios to the chromosome-level reference genomes of six *Stegodyphus* species from Ma et. al (2025) and a chromosome-level genome for *S. africanus* generated in this study (Figure S1, Table S1) to identify *de novo* mutations (DNMs) in the offspring.

All individuals were sequenced to a genome-wide mean depth of 27× (Table S2-3, Figure S2-4) after mapping quality control and genotype calling, and mutations were called according to the standard pipeline recommended by the Mutationathon (*53*). Quality filtering allowed us to genotype 165 trios (69 social, 96 subsocial, Figure 1D) out of 202 trios initially sequenced across all species (Table S4, See Supplemental Materials Section 3.7-3.8, Figure S5). We identified 712 *de novo* autosomal mutations (198 social, 514 subsocial, Table S5-6) and 41 *de novo* X-chromosome mutations (9 social, 32 subsocial, Table S7-8). There is restricted statistical power in differentiating the mutation rates of the X-chromosomes and the autosomes (Figure S6). The distribution of DNMs across the genome shows no clustering pattern in any species (Supplemental Methods and Materials Section 5.1, Figures S7-S14). All DNMs were manually checked in IGV (*54*) to remove false-positive calls. False-positive DNMs can also occur when sequencing depth is low in the parents, as heterozygous sites may be incorrectly called as homozygotes. We performed a sensitivity analysis of the depth criteria used for filtering, and the mutation rates detected were found to stabilise at depth 26; consequently, we used 26 as a minimum depth threshold (See Supplemental Methods and Materials, Figure S15).

**Figure 1.**
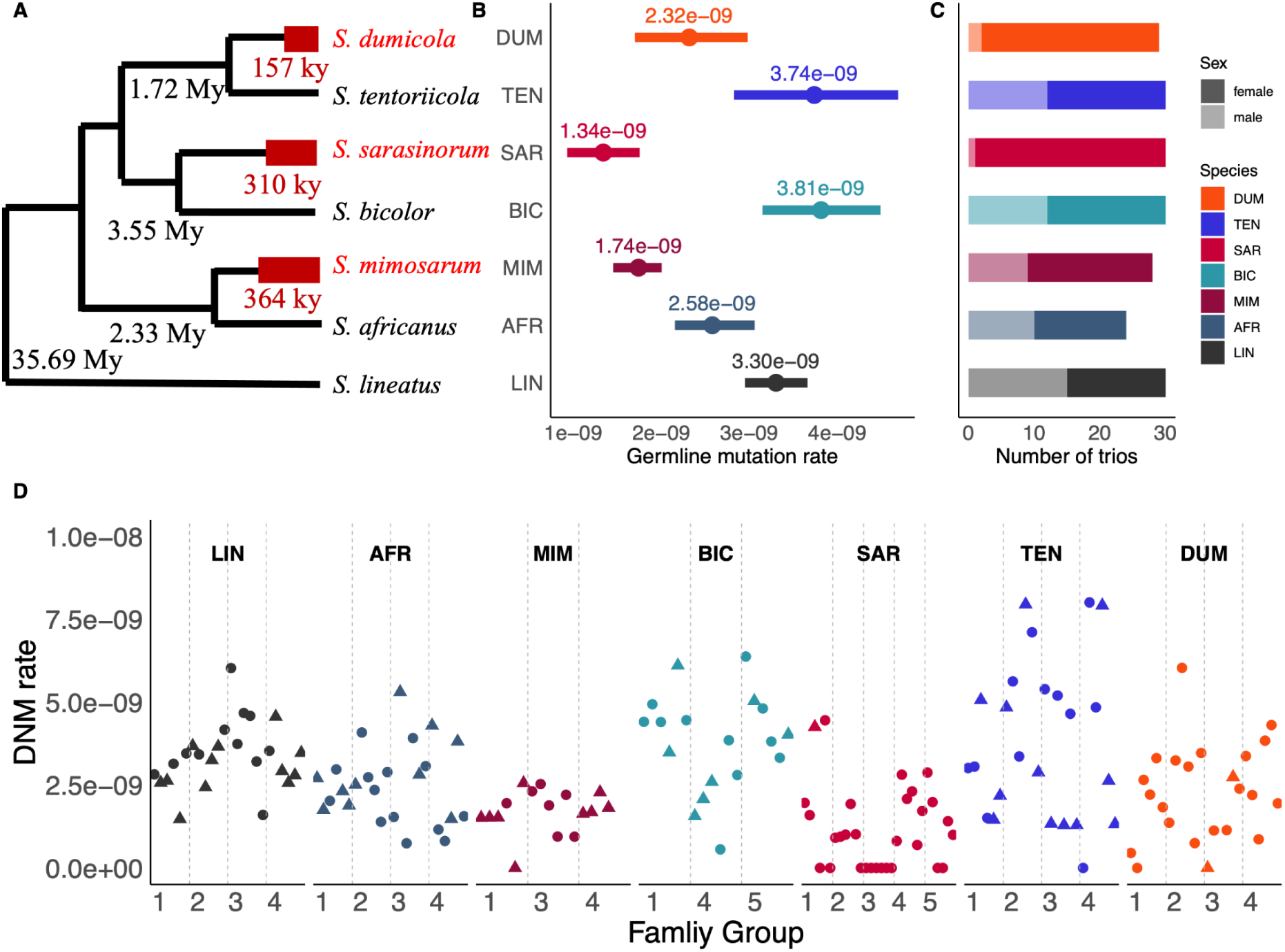
Germline mutation rate estimates in social and subsocial *Stegodyphus* spider species. (A) Phylogenetic tree with divergence time and estimated length of the social period (coloured in red) in seven species, recalibrated from Ma et al. 2025. The branch lengths in the figure are only illustrative and are not to scale. (B) The average germline mutation rate as estimated from trios for the seven *Stegodyphus* species. Two hypermutated families in *S. bicolor* are excluded from the species mutation rate estimation and following analyses (See Supplemental Materials and Methods, Figure S16). (C) The number of trios sequenced for each species and sex identification of offspring. Trios with female offspring are in solid colours, and trios with male offspring are in light colours. (D) Point estimation of germline mutation rates for all individuals that pass quality filtering (See Methods). Dots and triangles represent female and male offspring respectively. In *S. mimosarum*, the observed proportion of females was 67.9%, and not significantly different from 1:1 as in other social species, but showed high variation in sex ratios observed across each family: the female fractions of sequenced offsprings of the five families are 1/6, 6/6, 6/6, 0/4, 6/6 respectively.

We find that mutation rates are consistently lower in the three social species compared with the subsocial species (Figures 2B and 2D, Figures S16-18). The average germline mutation rate was 1.8-fold lower in social *Stegodyphus* species compared to the subsocial species (Figure 1B). The lowest estimated germline mutation rate was 1.34E-09 (bootstrapped 95CI: 0.93E-09 - 1.75E-09) in *S. sarasinorum,* and the highest germline mutation rate was 3.81E-09 (bootstrapped 95CI: 3.15E-09 - 4.49E-09) in *S.bicolor*. By contrasting mutation rate estimates in the three pairs of sister species in the phylogeny, the lineages leading to the social species exhibit lower mutation rates in all three comparisons.

**Figure 2.**
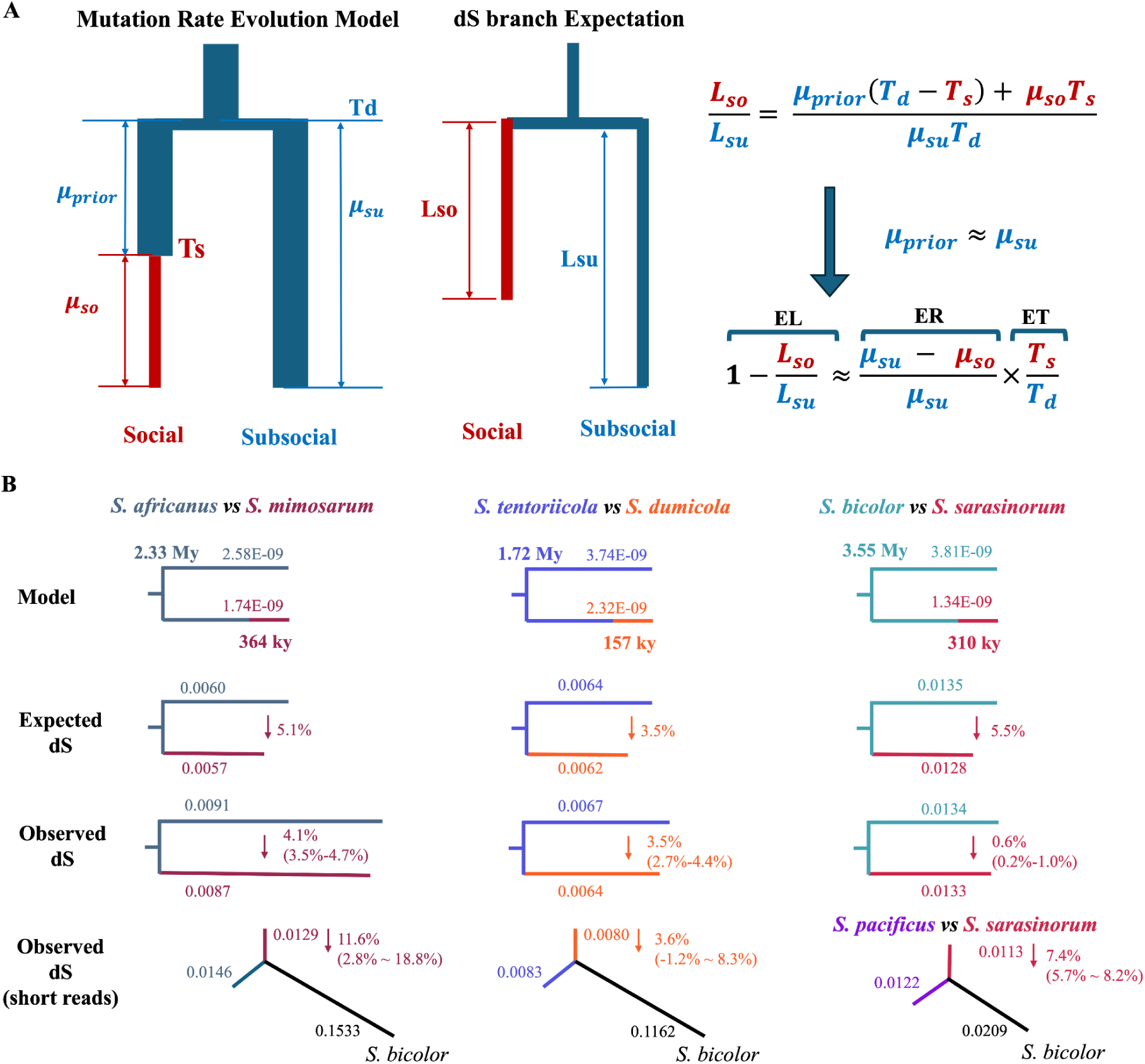
Mutation rate evolution and phylogenetic consequences. (A) Illustration of the mutation rate evolution model and the expected difference in *d_S_* branch lengths between pairs of social and subsocial spider lineages. The model assumes that mutation rate in the subsocial lineage represents the ancestral mutation rate, and that the germline mutation rate changes with the transition to sociality in the social lineage. EL denotes the reduction of the *d_S_* branch length (in percentage) in the social lineage. ER denotes the reduction of the mutation rate (in percentage) in the social lineage. ET denotes the fraction of the social period since the species divergence. (B) The parameters applied to the mutation rate evolution model, the expected *d_S_* branch length change and the observed *d_S_* branch length change in the three social-subsocial species pairs respectively. The branch lengths shown in the figure are illustrative for visualization.

In each social-subsocial species pair, the social species tended to show lower variation in individual mutation rate estimate compared to its subsocial sister species (Figure 1B and 1D, Supplementary Figure S19). However, the observed differences in mutation rate variation in all social-subsocial species pairs did not significantly deviate from uniform species-specific mutation rate simulations (Figure S20-21), suggesting that mutation rate variation does not systematically differ with degree of sociality.

### Phylogenetic evidence for reduced evolutionary rate in social lineages

If mutation rates decreased in the social lineages at the time of the transitions to sociality, then the terminal phylogenetic branches leading to social species should show a corresponding reduction in evolutionary rates (*55*). Under the neutral theory, the mutation rate equals the neutral substitution rate, suggesting that a reduction in mutation rate, as observed in the social species, should lead to a reduction in their synonymous branch length (*d_S_*). In all three comparisons, we see a reduction in the branch leading to the social species (Figure 2B, third row).

We modelled whether the reductions in *d_S_* are compatible with the estimated mutation rates determined here and the estimated timing of transitions to sociality from Ma et al (2025) (Figure 2A), assuming that (1) before the transitions, social lineages had the same mutation rate as those estimated in their subsocial sister species (Figure 1B), and (2) after the social transition, mutation rates declined instantly to the level estimated in the social species. We re-estimated social transition time and species divergence time from *d_S_*estimates in (*47*) using the mutation rate estimates from the present study (Figure 1B) instead of the mutation rate assumed in Ma et al (2025). This leads to the expected reduction in *d_S_* in each of the social branches shown in Figure 2B, second row (Table S10).

We performed two different estimations of the empirical reductions in *d_S_* to compare to those expected from the model; 1) branch-wise *d_S_* estimation from autosomal single-copy orthologs identified in Ma et al. (2025) based on long-read assembled reference genomes (Figure 2B, third row, Table S9), and 2) pair-wise *d_S_* estimation from the mapping of individual short-read resequencing data from pairs of species to a common outgroup, *S. bicolor* (Figure 2B, fourth row). The latter analysis allowed us to include *S. pacificus*, the closest subsocial sister lineage of *S. sarasinorum* even though it has not been long-read assembled (Figure 2B).

In the analysis based on assembled reference genomes, we find that the observed *d_S_* reductions match well with model predictions in *S. mimosarum* (4.1%; 95% CI, 3.5 to 4.7%; expected, 5.1%) and *S. dumicola* (3.5%; 95% CI, 2.7 to 4.4%; expected, 3.6%), but not in *S. sarasinorum* (0.6%; 95% CI, 0.2 to 1.0%; expected, 5.5%). The latter discrepancy could reflect the greater phylogenetic distance between *S. sarasinorum* and *S. bicolor* explained by the fact that *S. bicolor* is not the closest subsocial relative of *S. sarasinorum* (46–48). The short-read consensus analysis, which allowed a comparison between *S. sarasinorum* and the closest relative *S. pacificus*, found a 7.4% reduction in the *S. sarasinorum* lineage (95% CI, 5.7 to 8.2%; expected 9.4%, see Supplemental Section 6.3, Table S10), which is a close fit to the reduction expected under the model. Using short-read data mapped to *S. bicolor* as a reference further supported that dS is reduced in *S. dumicola* and *S. mimosarum*, although confidence intervals were much broader, likely due to increasing phylogenetic distance in the cross-species mapping (Figure S22).

We conclude that *d_S_* on branches leading to social species is consistent with our model of an instantaneous decline in mutation rate at the time of the transitions to sociality. This is probably a simplifying assumption of a complex transition process and we can not rule out that the reductions in mutation rates were more gradual over time (See Supplemental Section 6.5 and Figure S23). The results of the phylogenetic analyses of *d_S_* thus suggest that the low mutation rates measured in social species today are the end result of a reduction that occurred near the origin of sociality, rather than an environmental effect specific to the parents sampled in the trio experiments.

### Possible causes of reduced germline mutation rates in social species

We next investigated possible causes of reduced mutation rates in the social species by comparing in which parent and at which stage in germline development mutations occur, and by comparing their mutational spectra. Mutations that occur early in germline development will be shared among siblings (*2*). Contributions of DNMs are often sex-specific, with fathers contributing more DNMs to offspring than mothers in vertebrates (*5*). The specific types of mutations and their relative frequencies constitute the mutational spectrum, whose shape can reflect underlying mutagenic factors (*3*).

#### Absence of early mutations in social species

The 712 identified DNMs occurred at 676 unique chromosomal positions, as 27 DNMs were shared by two or more siblings. Strikingly, we found sibling-shared mutations only in subsocial species (Figure 3B, See Supplemental Materials Section 7.1, Figure S24-25). The shared mutations in subsocial species were 1% (1 out of 79, *S. tentoriicola*), 2% (2 out of 123, *S. africanus*), 4% (8 out of 173, *S. lineatus*), and 15% (16 out of 104, *S. bicolor*) of the total number of DNMs, respectively (Figure 3B). While fewer DNMs were detected in social species overall, the absence of identified shared mutations among siblings represents a statistically significant difference in the mutational process between social and subsocial species (Fisher’s exact test, p-value = 0.000114), supporting a contribution of a reduction of detectable early germline mutations in social lineages to the reduced mutation rate.

**Figure 3.**
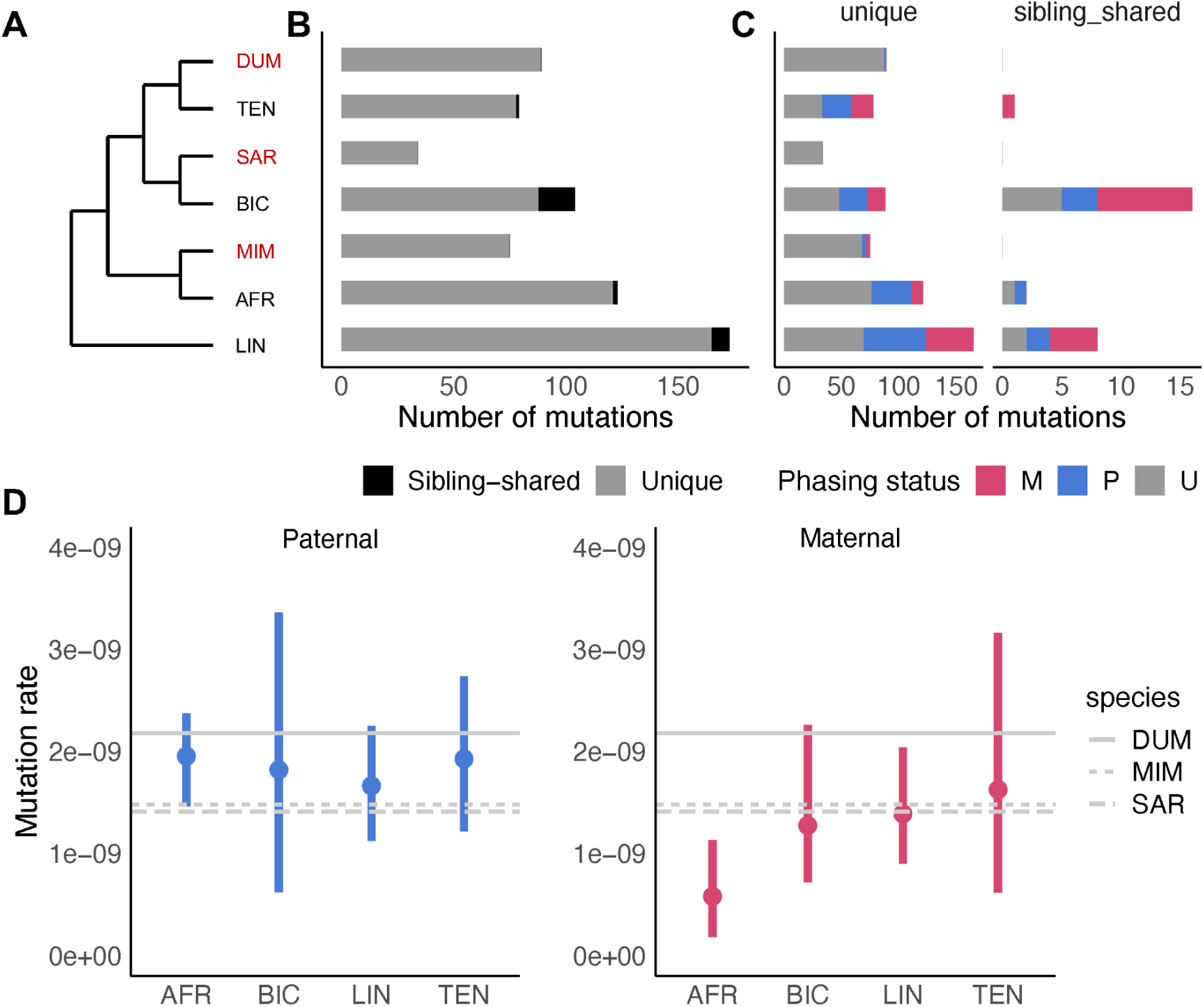
Paternal mutational differences between unique mutations and sibling-shared mutations. (A) Phylogenetic topology of the *Stegodyphus* genus, where social species are highlighted in red. (B) The number of detected mutations and status of sibling-sharing mutations across species. (C) The number of paternal mutations (P, blue), maternal mutations (M, red) and unphased mutations (U, grey) in the context of unique mutations and sibling-shared mutations. (D) The estimated paternal mutation rates (left, blue) and maternal mutation rates (right, red) across subsocial species, with comparisons to the germline mutation rates in social species (light grey horizontal lines).

We then used parental-origin phasing with manual validation to test whether early germline mutations were enriched for maternal or paternal origins (*53*, *56*, *57*). We succeeded in phasing 245 out of 479 DNMs (147 paternal and 98 maternal) in subsocial species (Figure 3C), but could only phase 9 out of 197 DNMs (6 paternal and 3 maternal) in social species due to the low heterozygosity (2.4% of that estimated in the subsocial species, Figure S26). In subsocial species (excluding *S. africanus*), unique DNMs showed an overall paternal excess of 1.41, which was strongest in the laboratory-raised *S. africanus* (parental excess of 3.5, Figures S27). In contrast, sibling-shared DNMs (18 out of 27 could be phased) were mainly of maternal origin: 13 were maternal and 5 were paternal, significantly different from the relative proportions among unique DNMs (Fisher’s exact P-value: 0.02273). Thus, the early germline mutations detected in subsocial species are disproportionately maternal.

The absence of early mutations in social species, together with the shift toward maternal bias in early mutations in subsocial species, suggests that the reduced mutation rate associated with sociality is partly due to a reduction in early germline mutations, probably linked to changes in oogenesis. This interpretation is consistent with the shift to producing fewer and larger eggs in females of social species (*39*). However, reduced early mutation rates alone are unlikely to explain the 1.8-fold reduction in the mutation rates of the social species. Even if all maternal mutations in subsocial species were assumed to be early germline mutations and completely absent in social species, the remaining paternal mutation rate in subsocial species would still exceed the total germline mutation rate observed in social species (Figure 3D). We therefore infer that the mutation rate reduction in social lineages most likely involves both maternal and paternal sources, with an additional contribution from a reduction in early-occurring DNMs.

#### The mutation rate of social species is reduced across the entire mutational spectrum

We next investigated whether the mutation rate in social species is reduced due to differences in the internal and external mutagenic pressures they experience. Such differences are likely to result in differences in their mutational spectra. We classified the DNMs into seven mutually exclusive classes (C>A, C>G, C>T (minus CpG > TpG), T>A, T>C, T>G and CpG>TpG) (Table S11). We first pooled DNMs from all seven *Stegodyphus* species and compared the mutational spectrum with those of the few other species for which more than 50 trios have been sequenced: sticklebacks, dogs, mice, and humans (Table S12). The overall shapes of the mutational spectra are similar across these taxonomically distant species and among the seven *Stegodyphus* species (Figure 4A-B, Figure S28), suggesting similarities in the mutagenic pressures across species and diverse taxonomic groups.

**Figure 4.**
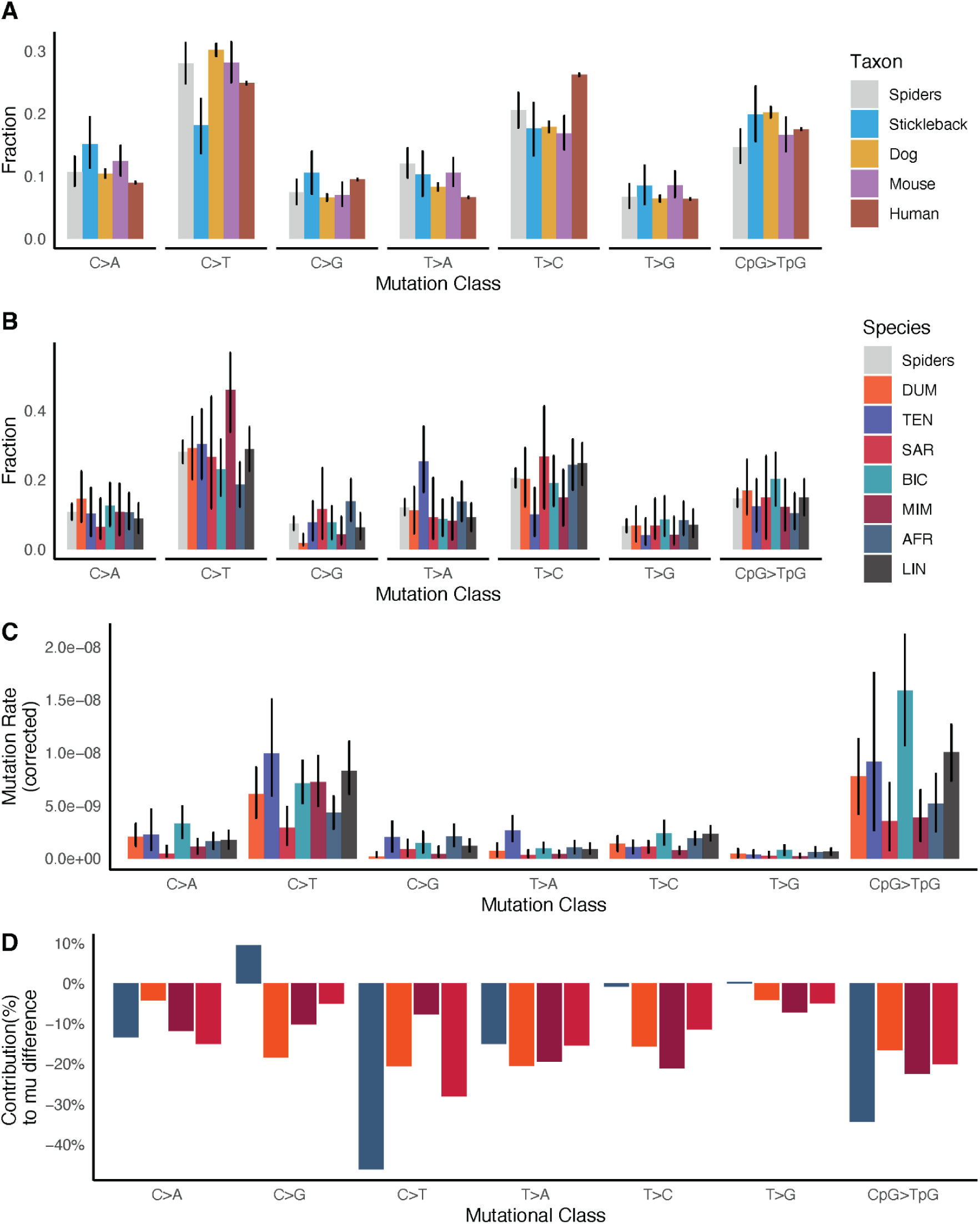
Germline mutational spectra of species in the *Stegodyphus* genus. (A) The variation of the mutational spectra in different taxa. (B) The variation in context-dependent mutation fraction among species of the *Stegodyphus* genus. The color code for species in the legend is shared from panel B to panel D (C) The germline mutation rates of different mutational classes in each species, after correction for the nucleotide composition of the callable genome per species. (D) The relative contribution to the observed mutation rate reduction from each mutational class in the lab-reared subsocial *S. africanus*, and the social species *S. dumicola*, *S. mimosarum* and *S. sarasinorum*. The mutation rate reduction in each species is evaluated as a comparison to the mean mutation rate of wild-caught subsocial species (*S. lineatus*, *S. tentoriicola*, and *S. bicolor*)

We calculated the mutation rate for each mutation class for each spider species (Figure 4C). In social species, the mutation rate is lower across the entire mutational spectra. We also calculated the average mutation rate for each mutation class for the three subsocial species with wild-caught parents and compared it to that of 1) the subsocial lab-raised *S. africanus*, and 2) the three social species that all have wild-caught parents, to evaluate the contribution of each mutational type to the observed lower mutation rates (Figure 4D, See Methods). For *S. africanus*, the reduction of the overall mutation rate is mainly driven by C>T mutations, both in and out of the CpG context. Several environmental mutagenic processes, such as UV-light and thermal stress, cause an increased rate in these types of mutations (*31*, *58–61*). Therefore, the relatively lower C>T mutation rates in *S. africanus* might be due to less exposure to UV-light and thermal stress in the lab. The lab environment is lit by artificial LED light without UV radiation and has a fluctuating temperature cycle between 21 and 29 degrees Celsius, both of which are less stressful than the natural environments.

In the social species, the reduction in mutation rate is spread relatively evenly across all seven mutation classes (Figure 4D). Combined with the generally stable germline mutational spectrum across species, this pattern indicates that the lower mutation rate in social species is unlikely to be caused by changes in specific mutagenic factors, such as reduced DNA damage from exposure to certain environmental mutagens. Instead, it is more consistent with a general slowdown in the accumulation of germline mutations. This raises the possibility that altered endogenous processes of germline development in social species underlie the reduction in mutation rate, although the exact mechanism remains unclear.

#### Somatic mutation and DNA repair gene expression

To further investigate the causes of the reduced germline mutation rate in social species, we examined somatic mutation rates and DNA repair gene activity in social and subsocial species. We used the PacBio HiFi reads generated for the reference genomes of individual species to determine the average somatic *de novo* mutation rate in each species, exploiting the high accuracy of HiFi sequencing (see Supplemental Methods and Materials Section 9). The somatic mutation rate was 3 to 15 times higher than the germline mutation rate across the species, as expected from mammalian somatic mutation rate studies (Figure 5B, Figure S29). Thus, even though we estimate the rate from a single individual, the number of *de novo* somatic mutations is sufficiently large (249 - 1679, Table S13, Figure S30) to obtain a precise estimate per individual. The results show that there is also a consistent reduction in somatic mutation rates in social species relative to their corresponding subsocial counterparts (Figure 5A-B). The parallel reduction in both germline and somatic mutation rates suggests that social species experience a broader decrease in mutation accumulation across tissues. This pattern points to two broad classes of explanations: First, mutation rates may be reduced due to life history and developmental changes associated with a transition to sociality. Fewer cell divisions (*5*, *62*, *63*) could lead to a lower mutation rate either by fewer replication errors or by more time to repair in more slowly dividing cells (*64*, *65*). This is plausible in social *Stegodyphus*, which despite similar development time are smaller in body size (Figure S31, Table S14) and produce fewer and larger eggs than their subsocial relatives (*39*), consistent with altered somatic growth and germline proliferation. Second, the reduction could reflect gradual molecular evolution toward generally improved DNA repair efficiency. Specifically, mutator alleles can be selected against more efficiently due to its increased linkage with deleterious mutations in a highly inbreeding system (*51*, *52*). Under this model, social species might show elevated expression of genes in DNA repair pathways (*66*), especially in germline-enriched tissues.

**Figure 5.**
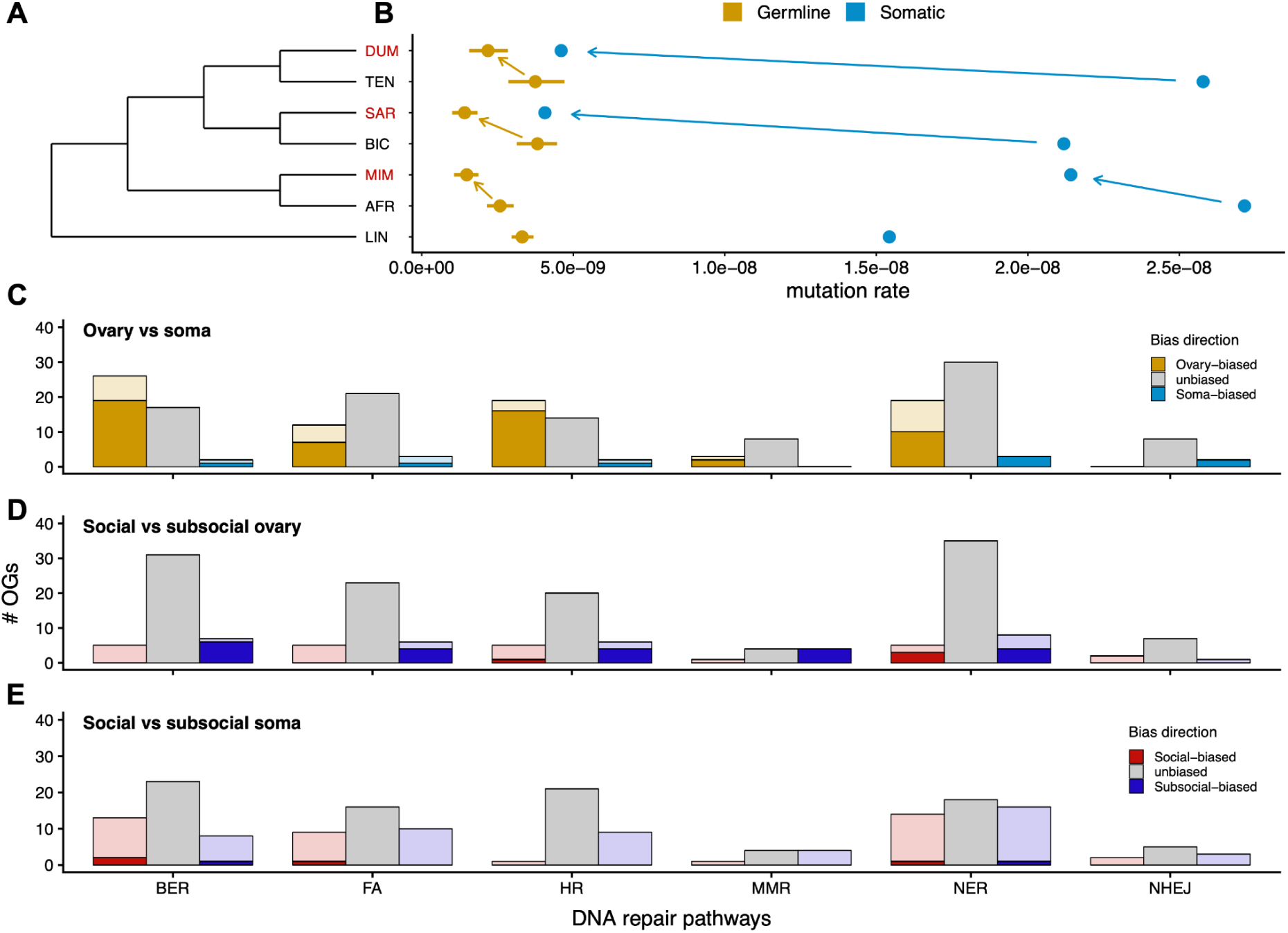
Differences in somatic and germline mutation rates across species and the corresponding differential gene expression levels in DNA repair pathways. (A) The phylogenetic relationship of the *Stegodyphus* genus. The phylogenetic tree is only to show the topology structure, not real branch lengths. (B) The somatic mutation rate and germline mutation rate estimated across all sampled species. The arrows denote the direction of mutation rate change from the subsocial species to the social species. (C) The number of differently expressed DNA repair genes between ovary tissue and whole body tissue and the direction of bias. A transparent color denotes the number of orthogroups with a more than 2-fold change between ovary and whole body tissue at the mean expression level across all species and the solid color denotes the fraction of orthogroups that consistently change in the same direction across all species. (D) The number of differentially expressed DNA repair genes between social spider species and subsocial spider species in the ovary tissue, where a transparent color denotes the number of orthogroups that are differentially expressed between social and subsocial species group and the solid color denotes the fraction of orthogroups whose bias directions fulfill the requirement of min(subsocial species means)>max(social species means) or vice versa. (E) The number of differentially expressed DNA repair genes between social and subsocial spider species in the whole body tissue, where a transparent colour denotes the number of orthogroups that are differentially expressed between social and subsocial species groups, and the solid colour denotes the fraction of orthogroups whose bias directions fulfill the requirement min(subsocial species means)>max(social species means) or vice versa. The abbreviations for pathways involved in DNA repair in the KEGG database are: base excision repair (BER), nucleotide excision repair (NER), mismatch repair (MMR), homologous recombination (HR), non-homologous end joining (NHEJ), and the Fanconi anemia (FA).

To test whether changes in mutation rates could be due to changes in the activity of DNA repair pathways, we analysed transcriptome data from ovary tissue in four species, including two social and two subsocial species (Figure S32), and adult whole-body transcriptomes from all seven species. Because germline tissues are generally more protected from mutations than somatic tissues, we first evaluated whether the expression of DNA repair pathways captures biologically meaningful differences between tissue types (Figure S33). Across KEGG pathways related to DNA repair, a substantial fraction of the total genes is differentially expressed between ovary tissue and whole body tissue (Figure 5C). Most of the differentially expressed genes had higher expression in ovary tissue than in adult whole-body tissue, and this pattern was broadly consistent across species (Figure 5C). Germline and somatic mutation rates and mutational spectra also differed significantly (Figure S29), consistent with distinct mutational processes between tissue types. Together, the higher DNA repair pathway expression in ovary tissue, the lower germline mutation rate, and the associated shift in mutational spectra supports DNA repair genes expression as an informative proxy for differences in tissue-level DNA repair capacity that are inversely related to mutation rates in *Stegodyphus*.

We then tested whether the lower mutation rates in social species could be explained by elevated DNA repair expression. In ovary tissue, most of the DNA repair genes are not differentially expressed between the social and subsocial species, and the differentially expressed genes were not biased toward higher expression in social species (Figure 5D, Figure S34). Instead, among genes showing consistent directional differences across all social-subsocial comparisons, more genes had higher relative expression in subsocial species than social species (Figure 5D). A similar pattern was observed in the adult whole body tissue (Figure 5E, Figure S35). Thus, we find no transcriptional signal for an upregulation of DNA repair pathways in social species. If DNA repair pathway expression is taken as an indirect proxy for repair efficiency, the observed direction in the ovary is opposite to what is expected under a scenario of enhanced DNA repair in social lineages.

## Conclusions

We show that the three independent transitions to sociality in *Stegodyphus* spiders are consistently associated with a pronounced, similar reduction in germline mutation rate. These lowered mutation rates occur despite the severe reductions in *N_e_* that accompany social evolution and are therefore unexpected under the drift-barrier hypothesis. Instead, we suggest that major life history transitions affect germline mutation rates. Reduced mutation rates occur across the mutational spectrum and lower mutation rates are also found in somatic tissues. However, the reduction is not accompanied by consistent changes in the transcriptional activity of DNA repair pathways. We therefore suggest that a slower life history with changes in developmental and reproductive traits, tightly associated with sociality, is the main reason for the reduced mutation rate.

Social lineages in *Stegodyphus* have been suggested to be evolutionary dead ends, potentially due to demographic vulnerability or genomic degeneration under severely reduced effective population size and selection efficacy. Our results indicate that their short evolutionary persistence is unlikely to be caused by an elevated germline mutation rate. Instead, the reduced germline mutation rate of social lineages may slow the accumulation of new deleterious single-nucleotide variants. Other processes may therefore better explain their short evolutionary persistence, including extreme female-biased sex ratios that increase demographic instability and sources of genomic degeneration other than single-nucleotide mutation, such as structural variants and transposable elements, under inefficient selection efficacy.

While the reduced mutation rate in social species is mirrored in a lower evolutionary rate on the phylogeny, we also observe that laboratory-raised *S. africanus* have lower rates of certain mutational types associated with environmental influences. This suggests that arthropod mutation rate estimates can be affected by the sampling environment (*67*), probably due to a more exposed germline than in e.g. mammals. Therefore, mutation rate estimates should consider the rearing environment as well as natural ecological contexts in arthropod species and other organisms with diverse germline and embryonic developmental processes.

## Acknowledgement

We thank Marie Braad Lund, Giulia Soffiantini for the sample collection and sample preprocessing. We thank Juraj Bergman, Brian Charlesworth and Deborah Charlesworth for many useful comments on the manuscript.

## Author contributions

Conceptualization: MHS, TB, JB, PV

Methodology: JM, MHS, JB, TB

Investigation: JM, JB, AA, PSP, AE

Visualization: JM, AA

Funding acquisition: TB, MHS

Data Processing: JM, AE, PSP, IK, SM, JN, MTPG

Supervision: MHS, TB, JB, PV

Writing – original draft: JM

Writing – review & editing: JM, MHS, TB, JB, AA, PA, PSP, PV

## Data and materials availability

All raw sequencing data in this project are submitted to NCBI Bioproject PRJNA994315 and PRJNA1234535.

The assembled chromosome-level *Stegodyphus africanus* genome assembly is submitted to NCBI under accession number GCA_056512425.1.

All custom processing scripts and processed metadata are uploaded to GitHub Repository https://github.com/Jilong-Jerome/Spider-de-novo-mutation-rate

