## Supplemental Methods and Materials for "Repeated Evolution of Sociality is Associated with Reduced Mutation Rates in Spiders"

### 1. Study design, sampling, and sequencing

##### 1.1 Species sampling and controlled crosses

Six of the seven *Stegodyphus* species were sampled from the same locations as those used for genome assembly by Ma et al. (2025). Subsocial species were sampled as subadults and brought to the laboratory at Aarhus University, where individuals were kept separately until sexual maturity. For each species, 8-10 male-female pairs were established by controlled mating. Males were frozen at -80 degrees C after mating. Mated females were kept individually until they produced egg sacs; after hatching, the mother and six offspring from each family were sampled and frozen at -80 degrees C.

*S. africanus* mated females that had not yet produced an egg sac were sampled in Pongola, South Africa, and brought to the laboratory to lay egg sacs. After hatching, offspring were reared to sexual maturity and separated as subadults. Controlled matings were then established between males and females from different clutches, following the same sampling procedure as for the other subsocial species, except that four families were obtained.

For each social species, five nests with subadult spiders were collected and brought to the laboratory. From each nest, multiple groups of six subadults were isolated in individual boxes, until the sexes could be identified. Only groups including five females and one male were kept. Approximately one week after all individuals were assessed to be sexually mature, the males were frozen at -80 degrees C and females were separated into individual boxes. Females that subsequently produced egg sacs were kept, and after the eggs hatched, the mother and six offspring were sampled and frozen at -80 degrees C. We aimed to analyze one family from each original nest, however, if not possible we sampled two families from the same original nest, and in two cases (both in *Stegodyphus mimosarum*) we sampled two families sharing the same father (Family2 and Family5, Family3 and Family4). After quality control, 165 trios were retained for de novo mutation analyses (Table S4).

#### 1.2 DNA extraction and trio short-read sequencing

DNA from the parents and six offspring from each trio were extracted using the Qiagen Blood and Tissue Kit and sequenced using short-read sequencing.

Whole-genome sequencing of 271 individuals (34 families, 65 parents (four families sharing the two fathers, one failed sequencing of a mother) and 206 offspring across 7 *Stegodyphus* species, Table S2-3) were performed on the same platform (DNBSEQ-G400). Sequencing was done in many small batches, with several individuals mixed per flowcell lane. Samples were distributed across 73 flowcells, with each carrying only a median of 4 individuals and ~2 families per flowcell, and individual families were spread across a median of 4 flowcells. Most flowcells contain sequencing from a single biological group (62 of 73 contained only one species and 33 only one family), and flowcell assignment was unstructured with respect to sociality (8 flowcells carried both social and subsocial individuals) and to parent/offspring (5 flowcells parent-only, 34 offspring-only, 34 carrying both; and in 20 of 34 families a parent and an offspring of the same trio were co-sequenced on a shared flowcell). Biological inference can consequently not be an artefact from batch effects. The complete per-individual mapping of species family, parent/offspring role, flowcell IDs, lanes and total coverage is given in Table S3 and visualized in Figure S4. The total depth of the raw sequencing reads were comparable across all individuals (Figure S3).

#### 1.3 PacBio HiFi sequencing for the *S. africanus* reference genome

DNA was extracted from a single *S. africanus* individual using the MagAttract HMW DNA Kit (Qiagen) according to the manufacturer's instructions. HiFi PacBio libraries were prepared using the SMRTbell prep kit 3.0. Fragment size was estimated to be approximately 7-11 kbp using a Femto Pulse system (Agilent) with the 165 kbp genomic DNA assay. HiFi libraries were bound to polymerase using PacBio Binding Kit 3.0 and loaded onto 10 PacBio 8M SMRT cells. Sequencing was performed on a Pacific Biosciences Sequel IIe instrument, producing 130 GB of fastq data for genome assembly.

##### 1.4 Hi-C library preparation and sequencing for the *S. africanus* reference genome

Two whole S. africanus specimens were pulverized, crosslinked, and processed using the Arima High Coverage Hi-C protocol for small tissues (Arima Genomics). Large proximally ligated DNA molecules were sheared by sonication using a Covaris LE220-plus (Covaris LLC) to an approximate fragment size of 500 bp. One library was prepared per specimen using the Arima Library Prep Module protocol. Libraries were sequenced by Novogene on an Illumina NovaSeq 6000 platform with 150 bp paired-end chemistry.

### 2. Reference genome resources

##### 2.1 Existing reference genomes

Reference genomes of *S. lineatus*, *S. tentoriicola*, *S. bicolor*, *S. mimosarum*, *S. dumicola*, and *S. sarasinorum* were retrieved from Ma et al. (2025).

##### 2.2 *S. africanus* genome assembly and annotation

The *S. africanus* reference genome was assembled and annotated using PacBio HiFi and Hi-C sequencing under the same pipeline as in Ma et al. (2025). The draft genome was assembled with hifiasm using PacBio HiFi and Hi-C reads together. Haplotype-resolved draft assemblies were further scaffolded using the 3D-DNA pipeline. The scaffolded longer haplotype assembly was reported as the reference genome.

The reference genome was annotated with BRAKER2 (*1*) using RNA-seq data from *S. africanus* from Bechsgaard et al. (2019) and protein homolog sequences from other *Stegodyphus* species. Assembly completeness was evaluated with BUSCO (*2*) using the OrthoDB v10 arthropod dataset (*3*) and with Merqury (*4*)(Table S1).

### 3. Germline de novo mutation discovery

##### 3.1 Read alignment and genotype calling

Reference genomes for each species were indexed with bwa-mem2 (*5*), and individual short-read data were aligned to the corresponding species reference genome. Samtools (*6*) and Picard were used to remove duplicated reads and retain only properly paired read alignments with mapping quality greater than 60. The remaining alignments were used for genotype calling in GATK4 (*7*).

GATK HaplotypeCaller was run with -ERC BP_RESOLUTION to obtain genome-wide gVCF files for each individual. GATK GenomicsDBImport was then used to build per-species databases including all individuals, followed by GATK GenotypeGVCFs with the -all-sites option to call genotypes across the genome, including non-variant sites. The recommended GATK germline variant filters were applied with VariantFiltration using the following thresholds: QD < 2.0, MQ < 40.0, FS > 60.0, SOR > 3.0, MQRankSum < -12.5, or ReadPosRankSum < -8.0.

##### 3.2 Individual depth summaries

Mean depth on autosomes and X chromosomes was calculated separately for each individual from the genome-wide VCF including all sites using vcftools (*8*). These individual chromosome-specific depth estimates were then used to define depth filters for callable genomic regions.

##### 3.3 Callable sites

Callable sites were defined per trio. For a chosen minimum-depth threshold (minDP: 20, 22, 24, 25, 26, 28, or 30), a site was considered callable in a trio only if all of the following criteria were met:

1. The genotype quality (GQ) was higher than 60 in all three individuals in the trio.
2. The depth (DP) was within 0.5x to 2x of the chromosome-specific mean depth for each individual in the trio.
3. The site was invariant or biallelic.
4. The depth was at least minDP; for male X chromosomes, the threshold was 0.5 x minDP.

##### 3.4 Candidate de novo mutation identification

Candidate DNMs were identified among callable sites by searching for Mendelian violations. For autosomes, a site was considered a putative DNM when both parents were homozygous for the reference allele and the offspring was heterozygous. For X chromosomes, both parents were required to carry the reference allele, and the offspring was required to be heterozygous in females or carry an alternative allele in males.

Additional candidate filters were applied in the following order:

1. The candidate was a single-nucleotide polymorphism.

2. No other Mendelian violation was observed within 150 bp in genotypes called with bcftools from the original BAM files.
3. Allelic balance at heterozygous sites was between 30% and 70%.
4. The candidate passed the GATK germline variant filters.

##### 3.5 Manual validation of DNM candidates

Candidate DNMs were checked using the original alignment BAM files with bcftools and manual inspection in IGV. Candidates were retained as high-confidence DNMs only when the heterozygous site in the offspring was supported by reads in the original BAM file and no parental reads carried the alternative nucleotide observed in the offspring.

##### 3.6 Final depth threshold and sensitivity analysis

Minimum-depth filters on callable sites were used to reduce false-positive DNM calls caused by unresolved parental heterozygosity. At low parental coverage, a truly heterozygous site, one of the alleles may by chance not be sequenced, and therefore genotyped as homozygous. If the unsampled parental allele is transmitted to the offspring and is observed as a heterozygous genotype, the site can appear as a Mendelian violation and be falsely classified as a DNM, leading to an upward bias in the mutation-rate estimate. This source of error is expected to decline as the minimum read depth required for callability increases.

To evaluate this effect, we repeated the callable-site definition and species-level germline mutation-rate estimation across a series of minimum-depth thresholds (minDP = 20, 22, 24, 25, 26, 28, and 30; with these thresholds being 50% applied to male X chromosomes, as described above). Estimated mutation rates decreased with increasing minDP at low thresholds, consistent with progressive removal of false positives due to hidden parental heterozygosity, but reached a plateau at minDP = 26 (Figure S15). We therefore used minDP = 26 for the reported mutation-rate estimates and all downstream analyses. This threshold balances conservative false-positive control with retention of callable genomic sites needed for power in species-level rate estimation and analyses of mutational processes.

##### 3.7 Kinship verification

To verify parent-offspring assignments and exclude labelling errors, we tested all trios with usable parental genotypes using a Mendelian consistency score that depends only on within-trio segregation and does not require population allele frequencies. For each putative trio, we considered autosomal biallelic SNPs that passed GATK filters and at which the two assigned parents were homozygous for opposite alleles (0/0 and 1/1). At these sites, a true offspring must be heterozygous; the consistency score was the fraction of informative sites at which the offspring genotype was 0/1. Correct trios are expected to have scores close to 1, apart from residual genotyping error, whereas misassigned parents produce a marked reduction.

Genotypes were taken from the per-species GATK joint call sets. Sex chromosomes were excluded because male hemizygosity violates the diploid Mendelian expectation. We applied no minor-allele-frequency filter, because informative sites are defined within each trio. Genotypes with $GQ < 20$ or $DP < 26$ were treated as missing, using the same depth threshold as in the DNM callability filter to minimize artefactual inconsistencies caused by allelic dropout.

After excluding families with parental sequencing failure (family 5 in *S. tentoriicola*, *S. mimosarum*, *S. lineatus*, and *S. africanus*), the test identified two families with incorrect kinship assignments: family 5 in *S. dumicola* and family 2 in *S. mimosarum* (Figure S5). These families were excluded from the DNM analysis. The test also confirmed that the father was shared between families 3 and 4 in *S. mimosarum*, making the offspring from these two families half-siblings.

##### 3.8 Hypermutated families

Two *S. bicolor* families, Family 2 and Family 3, had significantly higher mutation rate estimates than the remaining families in the species (Figure S16). Family 2 had a substantially smaller callable genome size, which increases uncertainty in its estimate (Figure S17). However, the two hypermutated families had very similar estimated mutation rates, and Family 3 had a callable genome size comparable to other *S. bicolor* families. Thus, the elevated rates were unlikely to be explained by low callability alone in Family 2.

The excess DNMs in these families were not restricted to a single mutation class. Both families showed elevated rates across mutational classes and a higher fraction of sibling-shared DNMs, suggesting that higher mutation rates are due to a general increase in mutation rate across all mutational classes rather than a technical artifact (Figure S28). Parental-origin phasing further suggested a maternal contribution in Family 3, where we identified 47 unique maternal DNMs and 11 unique paternal DNMs, opposite to the paternal bias observed across most other subsocial families (Figure S27).

Because the cause of these family-specific elevations cannot be resolved from the available data, we report them as exceptional cases and exclude them from species-level mutation-rate estimates and downstream evolutionary analyses.

### 4. Germline mutation rate estimation

**4.1 Autosomal mutation rate estimation**

For each species, we estimated the autosomal germline mutation rate by summing DNMs and callable sites across all retained trios; the species-level point estimate was calculated as the total number of autosomal DNMs divided by the total autosomal callable genome size. This estimator weights each trio by the number of callable sites it contributes, rather than giving every trio equal weight as would occur when averaging individual trio-level rates.

This weighting addresses the concern that low-callability families with zero observed DNMs could drive species rates downward. A trio with few callable sites has little opportunity to contribute DNMs, so observing zero DNMs can be expected even when the underlying mutation rate is similar to the species average. We therefore used the summed counts-and-callable-sites estimator for species-level point estimates and assessed uncertainty by resampling trios in the bootstrap analysis below.

**4.2 Bootstrap confidence intervals for autosomal rates**

Confidence intervals were estimated by bootstrapping trios with replacement within each species. For each bootstrap replicate, the number of sampled trios equaled the number of retained trios for that species. A mean mutation rate was estimated for each bootstrap replicate, and 95% confidence intervals were reported as the 2.5% and 97.5% quantiles across 1000 bootstrap replicates.

##### 4.3 X chromosome mutation rate estimation

Because power was limited for estimating X chromosome mutation rates by species, X chromosome rates were estimated for grouped categories: social species and subsocial species. *Stegodyphus africanus* was excluded from the subsocial category here for its being fully lab-reared. The average X chromosome mutation rate was calculated as the total number of X chromosome DNMs divided by the total callable X chromosome size of all individuals in the corresponding category. Female offspring contributed two sets of callable X chromosomes, whereas male offspring contributed one. Bootstrap confidence intervals were estimated using the same trio-level resampling approach as for autosomes.

#### 5. Quality-control and robustness analyses

##### 5.1 Genomic distribution of DNMs and clustering tests

To test whether DNMs were spatially clustered within genomes, we compared the observed average nearest-neighbor distance (ANND) among autosomal DNMs with a permutation-based null distribution. The test was performed independently for each of the seven *Stegodyphus* species. For each species, unique autosomal DNM events were pooled across all retained trios and grouped by chromosome, so the analysis evaluated species-level spatial structure rather than within-individual clustering.

For each observed dataset, the nearest-neighbor distance of a DNM was defined as the physical distance, in base pairs, to the closest other DNM on the same chromosome. The ANND was calculated as the mean of these distances across all DNMs with at least one same-chromosome neighbor; chromosomes with a single DNM did not contribute to the statistic. The same statistic and exclusion rule were applied to both the observed data and all permutations. Lower-than-expected ANND values indicate clustering, whereas higher-than-expected values indicate more even spacing than expected by chance.

Null distributions were generated by drawing N positions, where N was the number of unique autosomal DNM events for the species, uniformly from the empirical union of all callable autosomal segments. Only base pairs within this callable-genome union were eligible for sampling, and each eligible base pair had equal probability of being selected. For each species, ANND was recalculated for 1,000 permuted datasets.

Significance was assessed from the permutation distribution using one-tailed probabilities for clustering and overdispersion. The clustering p-value was the fraction of permutations with ANND less than or equal to the observed value, and the overdispersion p-value was the fraction with ANND greater than or equal to the observed value. The two-tailed p-value was calculated as twice the smaller of these probabilities. We also report a standardized effect size, Z = (ANNDobs - mean(ANNDperm)) / sd(ANNDperm), where negative values indicate clustering and positive values indicate overdispersion. Across all species, observed ANND values fell within the corresponding null distributions, with no significant departure from a uniform distribution across callable autosomal regions (all two-tailed $p > 0.05$, $|Z| <= 1.3$; Figure S14).

**5.2 Individual-level mutation rate variation**

To assess whether individual-level germline mutation-rate variability differs between social and subsocial *Stegodyphus* species, we calculated autosomal mutation rates for each retained offspring and compared species-level variability using Poisson simulations. Three families were excluded before this analysis: *S. bicolor* families 2 and 3, because they were classified as hypermutated, and *S. sarasinorum* family 3, because it had low callability and no observed DNMs.

For each offspring, the individual mutation rate was calculated as the number of unique autosomal DNMs divided by the autosomal callable-site. Within each species, individual-level variability was summarized as the coefficient of variation, CV = SD(r_i) / mean(r_i), using the population standard deviation and the corresponding inter-individual variance was also recorded. CV was used because it standardizes variability by the species mean and is therefore comparable among species with different average mutation rates.

We evaluated whether the observed CV exceeded that expected from count sampling alone under a species-specific Poisson null model. For each species, all offspring were assumed to share the summed species mean mutation rate. In each of 10,000 simulations, the DNM count for offspring i was drawn from Poisson(lambda_i = 2*C_i * mu_s), where C_i is its autosomal callable-site total and mu_s is the pooled species mean rate. Simulated counts were divided by C_i to obtain simulated individual rates, and the species CV was recalculated exactly as for the observed data.

Two social-versus-subsocial comparisons were then performed. First, we compared the mean CV across subsocial species with the mean CV across social species using ΔCV = mean CV(subsocial) - mean CV(social). The empirical one-tailed p-value was the fraction of simulations in which ΔCV was greater than or equal to the observed ΔCV, corresponding to a test for higher variability in subsocial species. Second, we repeated the comparison within the three phylogenetically matched social-subsocial pairs: *S. bicolor* with *S. sarasinorum*, *S. africanus* with *S. mimosarum*, and *S. tentoriicola* with *S. dumicola*. *S. lineatus* was excluded from the pairwise test because it has no social sister species in the dataset.

Individual-level CV varied approximately three-fold across species but showed no systematic separation by social system (Figure S19). The mean CV was 0.435 across subsocial species and 0.677 across social species. In the pooled comparison, the observed ΔCV was -0.242, close to the simulated null mean of -0.232, with a one-tailed p = 0.571 (Figure S20). The sister-pair comparisons gave the same conclusion: for each matched pair, the observed ΔCV fell within the Poisson null expectation (Figure S21). We therefore found no evidence that individual-level germline mutation-rate variability differs between social and subsocial *Stegodyphus* species and the observed differences are consistent with Poisson sampling noise.

### 6. Phylogenetic analyses of mutation-rate evolution

##### 6.1 Recalibration of species divergence and social-transition times

Species divergence times in Ma et al. (2025) were estimated by dividing mean $d_S$ branch length between species pairs by an assumed mutation rate of 5e-09 per site per generation and a generation time of one year. Social-transition times were estimated as a fraction of species divergence time for each pair of sister species.

Because the mutation-rate estimates in this study differ from the previous assumption, divergence and transition times were recalibrated by dividing the $d_S$ branch length of subsocial lineages by the average mutation rate across wild-caught subsocial species (*S. lineatus*, *S. tentoriicola*, and *S. bicolor*). *Stegodyphus africanus* was excluded from this analysis because its mutation rate may be reduced by the laboratory conditions and may therefore not reflect the rate acting over evolutionary history.

##### 6.2 Branch-wise $d_N$ and $d_S$ from chromosome-level assemblies

Bootstrap results for branch-wise $d_N$ and $d_S$ estimates in *Stegodyphus* were retrieved from Ma et al. (2025). The dataset contains 500 bootstrap replicates of $d_N$ and $d_S$ estimates based on alignments of 500 randomly sampled autosomal single-copy orthologs for each species lineage. For each bootstrap replicate, the reduction in $d_S$ in the social lineage relative to the corresponding subsocial lineage was calculated. Mean estimates were calculated across the 500 bootstrap replicates.

Confidence intervals were estimated by bootstrap replicates 1000 times for each species pair. The 2.5% and 97.5% quantiles of these bootstrapped mean estimates were reported as the 95% confidence interval.

##### 6.3 Expected $d_S$ reduction under a mutation-rate shift model

To evaluate whether the observed mutation-rate shifts could account for reductions in synonymous divergence ($d_S$), we used a simple proportional model. For each social-subsocial species pair, the expected $d_S$ reduction was calculated as the fraction of divergence time spent in the social state multiplied by the fractional reduction in the germline mutation rate of the social species relative to its subsocial comparator. Social-period fractions were taken from Ma et al. (2025), where they were inferred from branch-wise $d_N/d_S$ estimates and $pi_N/pi_S$ among divergent populations within species. These fractions were 9.16% for *S. dumicola-S. tentoriicola*, 8.46% for *S. sarasinorum-S. bicolor,* 18.78% for *S. sarasinorum-S. pacificus*, and 15.62% for *S. mimosarum-S. africanus* (Table S10). Using the point estimates from this study, the inferred mutation-rate reductions in social species were 37.97% for *S. dumicola* relative to *S. tentoriicola,* 64.83% for *S. sarasinorum* relative to *S. bicolor*, and 32.56% for *S. mimosarum r*elative to *S. africanus*. Multiplying these reductions by the corresponding social-period fractions gave expected $d_S$ reductions of 3.5%, 5.5%, and 5.1%, respectively (Table S10).

Because no direct germline mutation-rate estimate was available for *S. pacificus*, we evaluated a plausible 30-60% reduction in *S. sarasinorum* relative to *S. pacificus,* based on the range observed among the measured *Stegodyphus* comparisons. Given a social-period fraction of 18.78%, this corresponds to an expected $d_S$ reduction of 5.6-11.3%; the main text reports 9.4% under a 50% mutation-rate reduction.

Finally, the *S. africanus* trio parents were laboratory reared and the mutational spectrum analyses suggest that their reduced mutation rate may partly reflect environmental conditions. If the *S. mimosarum-S. africanus* comparison is instead calibrated using a 50% social-lineage mutation-rate reduction relative to a wild subsocial baseline, the expected $d_S$ reduction increases from 5.1% to 7.8%, closer to the observed short-read based $d_S$ reduction estimate.

#### 6.4 Short-read consensus $d_S$ analysis including *S. pacificus*

##### *6.4.1 Analysis overview*

*Stegodyphus pacificus* is the closest subsocial sister species of *S. sarasinorum,* but only short-read resequencing data are currently available for this species. To include *S. pacificus* in analyses of synonymous substitution rates, we generated short-read consensus genomes for all three social-subsocial comparisons using the *S. bicolor* reference genome as a common coordinate system and outgroup reference. For each comparison, reads from one representative individual of each query species were mapped to the *S. bicolor* assembly, genotyped, and converted into masked consensus genomes. Coding-sequence alignments were then extracted from the *S. bicolor* annotation and analysed with PAML codeml; uncertainty was estimated by gene-level bootstrap resampling.

##### *6.4.2 Read alignment and variant calling*

Paired-end reads from each query individual were aligned to the S. bicolor reference with bwa-mem2 mem v.2.2.1 using default parameters. Downstream processing used SAMtools v.1.17 and Picard v.2.25.4. Read groups were added with Picard AddOrReplaceReadGroups. Mate information was corrected and secondary or unmapped reads were removed with samtools fixmate -rm; alignments were then coordinate-sorted and PCR or optical duplicates were removed with samtools markdup -r. Final BAM files retained only properly paired mapped reads (samtools view flags -f 0x2 -F 0x4) with mapping quality >= 60 and were then indexed.

Variants were called separately for each query species from the final filtered BAM files with bcftools v.1.16. Genotype likelihoods were calculated with bcftools mpileup against the *S. bicolor* reference using a minimum base quality of 20 (-Q 20).

*6.4.3 Callable masks and consensus construction*

To identify regions with anomalous depth, per-base coverage was calculated from each final BAM file with bedtools genomecov -bga v.2.31.1. For each species, we calculated the coverage-weighted median depth across covered sites (depth > 0) and defined a callable-depth window as max(5, ceiling[0.5 x median depth]) to floor(2.0 x median depth). Intervals outside this range were merged into a species-specific uncallable-depth BED mask.

Biallelic SNPs were extracted with bcftools view, retaining sites with variant quality >= 20 and depth within the species-specific callable window. Each retained SNP was classified from the read pileup using pysam v.0.23.3 with minimum base quality 20. Homozygous-reference and homozygous-alternate genotypes were accepted from the genotype call. Heterozygous sites were labelled het_clean only when high-quality pileup bases consisted exclusively of the reference and alternate alleles, with both alleles supported by at least one read; heterozygous sites with a third allele or missing support for either expected allele were labelled het_unclean.

Consensus construction used explicit rules for resolving heterozygous sites. Under the strict policy, only het_clean sites were eligible for resolution and het_unclean sites were masked; under the relaxed policy, both het_clean and het_unclean sites could be resolved. The reported analyses used the strict random setting. In this setting, heterozygous alleles were chosen reproducibly by hashing a fixed seed (random_consensus_seed = 20260514) with the species name and genomic coordinates, making the allele choice deterministic but independent across sites.

For each query branch, homozygous-alternate sites and heterozygous sites resolved to the alternate allele were written to a selected-SNP VCF. Homozygous-reference sites, heterozygous sites resolved to the reference allele, and sites that could not be resolved under the active policy were not written to this VCF. Non-resolvable sites were instead added to a rejected-sites BED. The final mask for each query species was the union of its uncallable-depth mask and rejected-sites mask. Consensus genomes were generated with bcftools consensus by applying the selected-SNP VCF to the *S. bicolor* reference and replacing masked positions with N. The unmodified *S. bicolor* reference served as the third genome in each three-sequence comparison.

##### *6.4.4 CDS alignment and filtering*

Coding sequences were extracted from the *S. bicolor* reference and the two query-species consensus genomes using the *S. bicolor* gene annotation. For each gene, the transcript with the longest total CDS was selected. CDS exon coordinates were extracted, concatenated, and reverse-complemented for genes on the minus strand. Because all genomes were represented in the same *S. bicolor c*oordinate system, the resulting three-sequence alignments were positional and gap-free by construction. Non-ACGT characters were converted to N.

A gene was retained only if its CDS length was a multiple of three, the *S. bicolor* reference CDS contained no premature stop codon, and all three species had a callable, non-N fraction of at least 0.80. Passing genes were partitioned into autosomal genes (chromosomes bic_1-bic_14) and X-linked genes (chromosomes prefixed bic_X); the analyses reported here used autosomal genes only. Per-gene autosomal CDS alignments were concatenated into a supermatrix. The supermatrix was then filtered to retain only fully callable codons, requiring all three codon positions to be unambiguous A/C/G/T bases in all three species. The filtered codon supermatrix was written in PAML sequential PHYLIP format.

##### *6.4.5 Pairwise $d_N/d_S$ estimation and bootstrap confidence intervals*

Pairwise substitution rates were estimated from the filtered codon supermatrix with PAML codeml v.4.9 using runmode = -2 and model = 0. The analysis used codon sequences (seqtype = 1), the F3x4 codon-frequency model (CodonFreq = 2), the universal genetic code (icode = 0), estimated kappa (fix_kappa = 0, initial kappa = 2), and estimated omega (fix_omega = 0, initial omega = 1). Sites with ambiguity were removed by PAML cleandata = 1, complementing the upstream codon filter. $d_N/d_S$ was calculated only for comparisons with $d_S > 0$.

Point estimates of $d_N$, $d_S$, and $d_N/d_S$ were calculated from the full set of passing autosomal genes. Confidence intervals were estimated with 500 non-parametric bootstrap replicates, each generated by resampling passing genes with replacement to the same number of genes as the full dataset. For each replicate, the resampled genes were concatenated, codon-filtered, and analysed with the same codeml pairwise model. For each species pair, the bootstrap distribution was summarized by its mean, median, and 2.5th and 97.5th percentiles, with the latter defining the 95% confidence interval.

##### 6.5 Mutation rate reduction model and causal direction inference

An instantaneous mutation rate shift at the origin of sociality is a useful first-order approximation, but it is unlikely to capture all possible histories of mutation rate evolution. We therefore also considered a gradual mutation rate reduction model in which the rate declines over an interval around, or following, the social transition, without subsequent directional reversal. This model is biologically plausible because the traits that can influence mutation rate, including life history, reproductive schedule, physiology, and DNA-repair investment, may themselves change gradually during the evolution of sociality.

The gradual model is useful mainly as a sensitivity analysis. Under the simple non-reversible scenarios considered here, the expected reduction in accumulated $d_S$ is bounded between the instantaneous-shift expectation and approximately one half of that expectation, depending on whether the rate change begins at, or is centered on, the inferred social transition (Figure S23). Thus, the model broadens the range of expected $d_S$ reductions but does not add much power to confirm the association between sociality and lower mutation rate beyond the direct DNM estimates and comparative $d_S$ analyses. Fitting a more detailed gradual trajectory would require additional start, end, and shape parameters that are not identifiable from the present data. We therefore treat the gradual mutation-rate reduction model as an interpretive framework rather than an independent confirmatory test. It shows that a gradual history remains compatible with the observed $d_S$ patterns.

This limitation also constrains causal inference.

We see the following three explanations for the concurrent occurrence between social transitions and declines in mutation rate:

a. The transition to sociality itself reduces the mutation rate.
   Here, we specifically address the hypotheses that convergent life-history changes that are tightly linked to being social explain the reductions in germline mutation rate through slower oogenesis and development.
b. A lower mutation rate precedes (and possibly facilitates) the transition to sociality.
c. A trait that arises convergently and precedes sociality (for example, a reproductive change) causes a convergent reduction in the mutation rate.

To address the temporal order between social evolution and mutation rate reduction, we expanded our analyses of synonymous branch lengths and the mutation rate reduction mode (Figure S23) by including new sequencing data on the subsocial sister species *S. pacificus*. These analyses strengthen the inference that mutation rates declined near the origins of sociality in all three lineages, but we cannot distinguish whether the decline occurred simultaneously with, shortly before, or shortly after the transition to sociality. Thus, we are limited in resolving fully the timing and hence causality between transitions to sociality and the onset of reductions in mutation rate..

Based on available evidence, scenario (b), in which a low mutation rate preceded and enabled sociality, is less likely. If reductions in mutation rate would precede sociality, we would expect the closely related subsocial lineages to show similarly low mutation rates as those observed in social lineages. However, we observe the opposite pattern: the subsocial species consistently have substantially higher mutation rates than the social species.

In contrast, scenarios (a) or (c) are more plausible. First, the phylogenetic analysis of synonymous branch lengths suggests that convergent reductions in mutation rate are tightly associated with the origin of social transitions. Second, the consistent shifts in reproductive traits of social spiders (fewer, larger eggs) and developmental traits (smaller body size) can be linked to alterations in the mutational process. This interpretation is further supported by the finding of declines in early germline mutation rate and a simultaneous reduction in somatic mutation rate in social species.

Based on these lines of evidence, we propose that the transition to sociality is associated with life-history changes that drive mutation reductions in social lineages. We acknowledge that this interpretation relies on an assumed direct causal relationship between the transition to sociality and reductions in mutation rate derived from our data and analysis.

### 7. Shared mutations and parental-origin analyses

##### 7.1 Identification of shared DNMs

Shared DNMs were first identified within each species as DNM calls occurring at the same chromosome and coordinate in more than one offspring. We then used family labels and the kinship-verification results to distinguish DNMs shared among full siblings within a retained family from position-shared DNMs involving offspring assigned to different families.

Only one cross-family shared event was detected. This DNM occurred in two *S. mimosarum* offspring assigned to Family 3 and Family 4 at mim_14:70086473. Manual IGV inspection supported the DNM call in both offspring and did not indicate a false positive (Figures S24 and S25). Although the reads did not resolve the parental origin of the mutation, kinship verification showed that Families 3 and 4 shared the same sire. The most parsimonious explanation is therefore a genuine early paternal germline mutation, or paternal mosaic mutation, transmitted through the shared sire to two half-sibling offspring.

We did not include this event in the subsequent analyses of sibling-shared DNMs. Those analyses were designed to quantify sharing among full siblings within the same retained family, where shared DNMs have a consistent pedigree interpretation and sampling unit. The S. mimosarum event is a single half-sibling, cross-family case with a different pedigree structure and no comparable replication across species or families. Including it would mix distinct forms of mutation sharing and give disproportionate weight to one exceptional pedigree. We therefore report this event separately as a likely genuine early mutation but exclude it from downstream sibling-shared DNM summaries.

##### 7.2 Manual phasing of parental origin

The parental origin of DNMs was phased manually by inspecting IGV alignments in a 300 bp region around each DNM. A DNM was assigned to the maternal or paternal haplotype when it was in complete read-backed linkage with one or more variants unique to either the mother or the father in the trio. Linkage was confirmed when the DNM and parental-informative variant were observed together on the same read.

Because parental-origin assignment of DNMs depends on direct linkage between the de novo allele and nearby phase-informative inherited variants, we used conservative read-backed IGV phasing rather than statistical haplotype phasing; this avoids relying on population-based phase inference for singleton DNMs (*9*, *10*) and prevents differential phasing power and preciseness between high-heterozygosity subsocial species and low-heterozygosity social species from biasing comparisons.

##### 7.3 Estimation of paternal and maternal mutation rates from phased DNMs

For each family, the fraction of phased DNMs assigned to paternal or maternal origin was used to assign unphased DNMs probabilistically to the paternal or maternal class. Maternal and paternal mutation rates were then calculated as the number of phased plus assigned mutations from each parental source divided by the total family-level callable size. Assignment of unphased mutations was repeated 1000 times to obtain bootstrapped family-level paternal and maternal mutation rates. Species-level estimates were calculated by averaging the bootstrapped family-level estimates, with 95% confidence intervals from the 2.5% and 97.5% quantiles.

##### 7.4 Statistical tests of sibling-shared DNMs and parental-origin bias

Using autosomal germline DNMs, we tested two features of mutation sharing among full siblings. First, we asked whether the proportion of sibling-shared DNM loci differed between subsocial and social species. Second, among the subsocial species with sufficient phased sibling-shared DNMs, we tested whether sibling-shared and unique mutations differed in parental origin.

For the sociality comparison, each retained DNM locus was classified as unique or sibling-shared and pooled by sociality after excluding BIC_family2, BIC_family3, and SAR_family3. This produced the following 2 x 2 contingency table.

| Sociality | Sibling-shared loci | Unique loci | Total |
|---|---|---|---|
| Subsocial (AFR, BIC, LIN, TEN) | 27 | 452 | 479 |
| Social (DUM, MIM, SAR) | 0 | 198 | 198 |

We tested the association between sociality and mutation-sharing class using a two-sided Fisher's exact test (scipy.stats.fisher_exact). Sibling-shared loci were detected only in the subsocial group (27/479 loci, 5.6%) and were absent from the social group (0/198 loci), giving a significant excess of sharing in subsocial species ($p = 1.1 \times 10^{-4}$). Because one cell contained zero, the uncorrected odds ratio is undefined; after adding a pseudocount of 1 to each cell, the odds ratio was 12.3.

For the parental-origin comparison, we restricted the analysis to the subsocial species with sufficient phased sibling-shared mutations: *S. bicolor*, *S. lineatus*, and *S. tentoriicola*. Hypermutated families BIC_family2 and BIC_family3 were excluded. *S. africanus* was not included because all individuals were lab-reared and this species showed distinct mutation-rate, mutational-spectrum and paternal bias patterns relative to the other subsocial species. Phased DNM loci were collapsed by locus, classified as unique or sibling-shared, and pooled across the three species to produce the following contingency table.

| Mutation class | Paternal | Maternal | P/M ratio |
|---|---|---|---|
| Unique | 106 | 75 | 1.41 |
| Sibling-shared | 5 | 13 | 0.38 |

We tested parental-origin differences between unique and sibling-shared loci with a two-sided Fisher's exact test on this 2 x 2 table. Unique mutations were paternally biased (P/M = 1.41), whereas sibling-shared mutations were maternally biased (P/M = 0.38). The contrast was significant (odds ratio = 3.67, p = 0.023).

### 8. Mutational spectrum analyses

##### 8.1 Classification of DNM classes

DNMs were classified into seven mutually exclusive mutation classes after collapsing complementary strands: C>A, C>T, C>G, T>A, T>C, T>G, and CpG>TpG. Shared mutations were counted only once when estimating the mutational spectrum because they are expected to derive from a single mutation event.

##### 8.2 Mutational spectrum estimation

Mutational spectra were calculated across scenarios defined by species, sociality, and whether a mutation was shared among siblings. For each scenario, the mutational spectrum was calculated as the fraction of DNMs in each mutation class, such that fractions summed to one.

##### 8.3 Callable-site denominators for mutation-class rates

Callable sites for each trio were classified as A/T sites, C/G sites, or CpG dinucleotide sites. Mutation rates for each mutation class were calculated using the corresponding callable-site denominator. Mean rate estimates and 95% confidence intervals were obtained by 1000 rounds of trio-level bootstrapping within each species.

##### 8.4 Contribution of mutation classes to observed rate reductions

Mutation-class contributions to mutation-rate reductions were estimated for cases with reduced mutation rates relative to wild-caught subsocial species: S. africanus and the three social species. The average mutation rate of the wild-caught subsocial species was used as the reference. For each mutation class and case species or group, the contribution to the total reduction was calculated from class-specific mutation rates after accounting for the genomic fraction of the corresponding nucleotide context.

The total mutation rate change observed between each of the case (j) with a lower mutation rate compared to the average mutation rate in wild-caught subsocial species (mean) can be written as the following, where μ denotes the mutation rate, and f denotes the fraction of the corresponding nucleotide in the genome. $i \in \{C > A, C > T, C > G, T > A, T > C, T > G, CpG > TpG\}$

$j \in \{S.\ mimosarum,\ S.\ dumicola,\ S.\ sarasinorum,\ S.\ africanus\}$

$$\Delta\mu_{i,j} = \mu_{i,j} - \mu_{i,mean}$$

$$\Delta\mu_{j} = \sum_{i} \Delta\mu_{i,j} \times f_{i,j}$$

Then the contribution ($c_{i,j}$) of each mutation type (i) to the observed reduction in each case (j) can be represented by the following.

$$c_{i,j} = \Delta\mu_{i,j} / \Delta\mu_{j}$$

$$\sum_{i} c_{i,j} = 1$$

### 9. Somatic mutation analyses

##### 9.1 Somatic tissue sampling

PacBio HiFi reads used for somatic mutation analyses were obtained from Ma et al. (2025) for *S. dumicola*, *S. tentoriicola*, *S. mimosarum*, *S. bicolor*, *S. sarasinorum*, and *S. lineatus*. The *S. africanus* HiFi dataset was generated in this study.

##### 9.2 Somatic mutation calling

For each species, PacBio HiFi reads were assembled de novo with hifiasm v0.16.1 in default diploid mode. The gzipped FASTQ file for each sample was provided as the sole input. Hifiasm uses phasing information in the HiFi reads to generate two haplotype-specific assembly graphs, haplotype 1 (`*.bp.hap1.p_ctg.gfa`) and haplotype 2 (`*.bp.hap2.p_ctg.gfa`), together with a partially phased primary contig graph (`*.bp.p_ctg.gfa`). Each GFA was converted to FASTA by extracting segment

(`S`) records, using the segment identifier as the FASTA header and the segment sequence as the contig sequence.

This produced three FASTA references per sample: haplotype 1 (`{species}_hap1.fa`), haplotype 2 (`{species}_hap2.fa`), and primary contigs (`{species}_p_ctg.fa`). The two haplotype assemblies were also concatenated to form a combined diploid reference (`{species}_combined.fa`). The same HiFi reads used for assembly were aligned back to each reference with minimap2 v2.30 using the `map-hifi` preset. Alignments were streamed to SAMtools v1.22.1, converted to BAM, coordinate-sorted, and indexed. For each sample, reads were mapped independently to the combined diploid, haplotype 1, haplotype 2, and primary-contig references, producing four sorted and indexed BAM files.

Somatic mutations were called with an in-house pipeline designed for diploid PacBio HiFi data. The pipeline used the BAM file from mapping the HiFi reads to the built combined diploid assembly.. It scanned the BAM for high-confidence mismatches, requiring mismatch base quality >= 93, neighbouring base quality >= 45, and no indel within 10 bp of the mismatch. Mismatches overlapping low-complexity regions were excluded.

Candidate mutations were then validated against the haplotype-specific de novo assemblies. A candidate was retained only if the same read carried the same mismatch when aligned to the haploid assembly, supporting the mutation as a true somatic event rather than a mapping or assembly artefact. Finally, mutations within 1500 bp of either read end were excluded because PacBio HiFi reads show elevated false-positive rates near read termini (*11*).

Reported mutations were strand-collapsed to a pyrimidine reference base. Calls originally referenced to a purine (A or G), together with their flanking trinucleotide context, were reverse-complemented so that every mutation was represented with a C or T central base.

##### 9.3 Somatic callable sites and mutation-rate estimation

We estimated species-level per-site somatic single-nucleotide variant (SNV) rates using called somatic SNVs as the numerator and matched callable trinucleotide opportunities as the denominator. Rates were calculated for all SNVs combined and for seven mutation classes: C>T, C>G, C>T at non-CpG sites, C>T at CpG sites, T>A, T>C, and T>G.

Callable opportunities were counted directly from the raw PacBio HiFi reads for each species, rather than from the reference genome, so that the denominator matched the sequenced molecules screened for somatic mutations. For each read of length L, we excluded T = 1500 bp from both ends, matching the read-edge filter used during somatic variant calling. We then counted every overlapping trinucleotide whose central base fell within the retained interval [T, L - T). A counted trinucleotide could include one flanking base from the trimmed boundary, but trimmed positions were never counted as central bases. Trinucleotides containing non-ACGT bases were skipped, and reads with L <= 2T contributed no callable windows.

For each mutation class c, we constructed a class-specific denominator $D_c$ from the 32 canonical pyrimidine-centered trinucleotide counts T(xyz), where x and z are flanking bases and y is the central pyrimidine. The somatic mutation rate for class c was then calculated as $r_c = n_c / D_c$, where $n_c$ is the number of called somatic mutations assigned to that class. This procedure ensured that numerator and denominator were defined over the same per-read intervals and mutation-context classes.

### 10. RNA sequencing and DNA repair pathway analyses

##### 10.1 Whole-body RNA-seq sampling

For six of the seven species, individuals were sampled for whole body somatic tissue RNA sequencing as described in Ma et al (2025), while the seventh species, *S. africanus*, was sampled from South Africa and brought to the lab. For all species, three to five families were established by controlled matings, and 3 spiderlings, 3 subadults and 3 adults were sampled for RNA sequencing. For details on the establishment of families, see supplementary materials in Ma et al (2025). RNA was extracted from all individuals using Qiagen RNeasy Mini kit, and libraries were built using NEBNext Ultra II Directional RNA Library Prep Kit. Individual sequencing was done on an Illumina NoveSeq 6000 with the aim of obtaining ~6GB 150PE data.

#### 10.2 Reproductive-tissue RNA-seq sampling

Four species were sampled for RNA sequencing of ovary tissues, two social species: *S. mimosarum* and *S. dumicola* and two subsocial species: *S. bicolor* and *S. africanus*. Ovary dissections were done for 6-10 females of each species, and the best three samples, evaluated by eye, were chosen for RNA extraction (Figure S31). Ovaries were dissected in cold 1× EBR (10mM HEPES pH7.3, 130mM NaCl, 5mM KCl, 2mM CaCl2) on ice. Samples were homogenized with a plastic pestle and lysed in 1 mL of TRIzol. Chloroform (200 µL) was added, and samples were vigorously shaken, incubated for 3 min at room temperature, and centrifuged at 12,000g for 15 min at 4°C. The RNA-containing phase was transferred to a clean RNase-free tube with 1 µL of Glycogen coprecipitant, 550 µL of isopropanol was added, and samples were centrifuged at 16,000g for 30 min at 4°C. Supernatant was discarded from the RNA pellet and 1 mL of ice-cold 75% ethanol was added. Samples were centrifuged at full speed for 5 min at 4°C, and the supernatant was removed. Samples were briefly centrifuged at full speed at 4°C and residual ethanol was removed from the RNA pellet. Samples were dissolved in 15 µL of RNase-free water.

#### 10.3 RNA-seq read processing and quantification

RNA-seq reads were quality assessed with FastQC v. 0.12.1 and MultiQC v. 1.26 (*12*). Adapters and low-quality bases were trimmed with Trimmomatic v. 0.39 (*13*) using ILLUMINACLIP:TruSeq3-PE.fa:2:30:10, a sliding-window filter of 4:15, and a minimum retained read length of 30 bp. Genomes were indexed with STAR v. 2.7.11b (*14*) using a splice-junction overhang of 149 and GFF3 information passed with --sjdbGTFtagExonParentTranscript Parent and --sjdbGTFtagExonParentGene ID. Trimmed reads were aligned to the corresponding species reference genome with STAR. SAM files were indexed with SAMtools v. 1.21 (*15*) and quality checked using Qualimap (*16*) RNA-seq mode v. 2.3, SAMtools, and MultiQC. Before read counting, annotation GFF3 files were converted to GTF using agat_convert_sp_gff2gtf.pl from AGAT v. 1.4.2 (*17*) with --gtf_version 3. Reads were quantified at the gene level using featureCounts from Subread v. 2.0.8 (*18*) with splice-junction inference enabled.

#### 10.4 Orthogroup and functional annotation

GO terms and KEGG pathways were assigned to the genes from each species using eggNOG-mapper v. 2.1.12 (*19*) with DIAMOND (*20*) search database v. 2.1.12 and the settings --target_orthologs all, --go_evidence all, --pfam_realign none, and --report_orthologs. To complement eggNOG annotation, protein sequences were searched against the FlyBase *Drosophila melanogaster* proteome (*21*) using NCBI BLAST+ v.2.16.0 (*22*) with an E-value threshold of 1e-05 and a maximum of five target sequences retained per query.

#### 10.5 Differential expression between reproductive and somatic tissues

Repair gene differential expression between reproductive and adult somatic tissue was done for the subset of species for which reproductive ovary tissue was available: *S. africanus*, *S. bicolor*, *S. dumicola* and *S. mimosarum*. The cross tissue analyses were run on each species separately, but for cross species comparison purposes, paralogs were handled by summing the counts for each orthogroup in each sample. We subsequently removed orthogroups with expression counts < 10 across all samples and fitted a DESeq2 v. 1.46.0 (*23*) model using tissue type as a single factor. The contrast was ovary vs adult and thus positive log2 fold-change indicates higher expression in reproductive tissues for these analyses.

The genes involved in KEGG repair pathways, namely base excision repair (BER), nucleotide excision repair (NER), mismatch repair (MMR), homologous recombination (HR), non-homologous end joining (NHEJ), and the Fanconi anemia (FA) pathway, were extracted. Summary statistics were computed as the total number of genes within each KEGG pathway as well as a measure indicating the mean log2FC between tissues across species and the consistency across species: 1) the number of genes that show an average log2FC > 1 between ovary tissue and whole body tissue across all four species. 2) the number of genes where the direction of log2FC was consistent across all four species.

##### 10.6 Differential expression between social and subsocial species

Differential expression analyses using were done cross species using limma (voom) v.3.62.1 (*24*) in R v. 4.4.2 between social and subsocial species for A) all species adult somatic tissue - 3 social and 4 subsocial species and B) four species where ovary data is available - 2 socials and 2 subsocial species. In these two analyses paralogs were handled in three different ways, in order to also take gene length variation between species into account All three strategies yielded similar summary results, and only one is shown in the results. 1) “RPK rounded”, where raw expression counts are divided by gene length in kb, summed for each orthogroup and subsequently rounded to the nearest integer. These data were then run through limma-voom for DE analysis. This is the analysis presented in the results. 2) “RPK float” is similar to “RPK rounded”, but kept as floats, then passed to voom for DE analysis. 3) “Raw+sumLen”, where raw integer expression counts were summed per orthogroup and gene length (log2 of total orthogroup length in kb) was passed as an offset to the DE analysis in voom. For all three strategies, a TMM normalization factor was estimated per sample using edgeR v. 4.4.0 (*25*) and voom precision weights were computed to account for the mean-variance relationship. In cross species analyses, some orthogroups may not be present across species, imposing issues when calculating Voom precision weights, since it expects counts and does not handle NA values. Therefore the precision weights are calculated using structural zeros, that are then converted to NA before running the DE model. To evaluate whether results are consistent with a more conservative approach, we also did the precision weight calculation using only 1:1 orthogroups across species. Only results from the structural zero method are shown in results. A linear model was fitted using the design: ~ social type + species, where species were included as a blocking variable to account for technical baseline differences. The p-value of the model results were shrunk using eBayes shrinkage with robust=TRUE. The comparison is social vs. subsocial and thus positive log2FC is interpreted as more expressed in social species. We also fitted a model with social type as a random effect using limma duplicateCorrelation(block = species), and this analysis did not change the direction of the results. Repair pathways were extracted and summary statistics were calculated for each subanalysis as follows: number of genes for each repair pathway, and a number indicating consistency which included genes where all species showed a logFC in the same direction as the DE-analysis result (min(social species means) > max(subsocial species means) or vise versa).

### 11. Supplemental tables

Table S1. *S. africanus* genome assembly metrics.

Table S2. The sequencing genotype coverage called per individual.

Table S3. Per-individual sequencing batch, lane, family, role, and raw coverage metadata.

Table S4. Trio structure and sample metadata.

Table S5. Autosomal DNMs.

Table S6. Autosomal callable-site summary per trio.

Table S7. X chromosome DNMs.

Table S8. X chromosome callable-site summary.

Table S9. $d_S$ branch length observations from Ma et.al 2025.

Table S10. $d_S$ branch length expectations.

Table S11. Autosomal DNM mutational type classification.

Table S12. DNM mutational spectra across species.

Table S13. The somatic mutations and callable sizes identified across species.

Table S14. The body sizes in *Stegodyphus*.

#### 12. Supplemental figures

Following are the caption descriptions and the corresponding figures embedded. All figures in PDF or PNG format in high resolution can be found both in the supplemental Data S1 and https://drive.google.com/drive/folders/1rWJ807kWhxTj9qfGbFSPNqMriYcKifef?usp=sharing

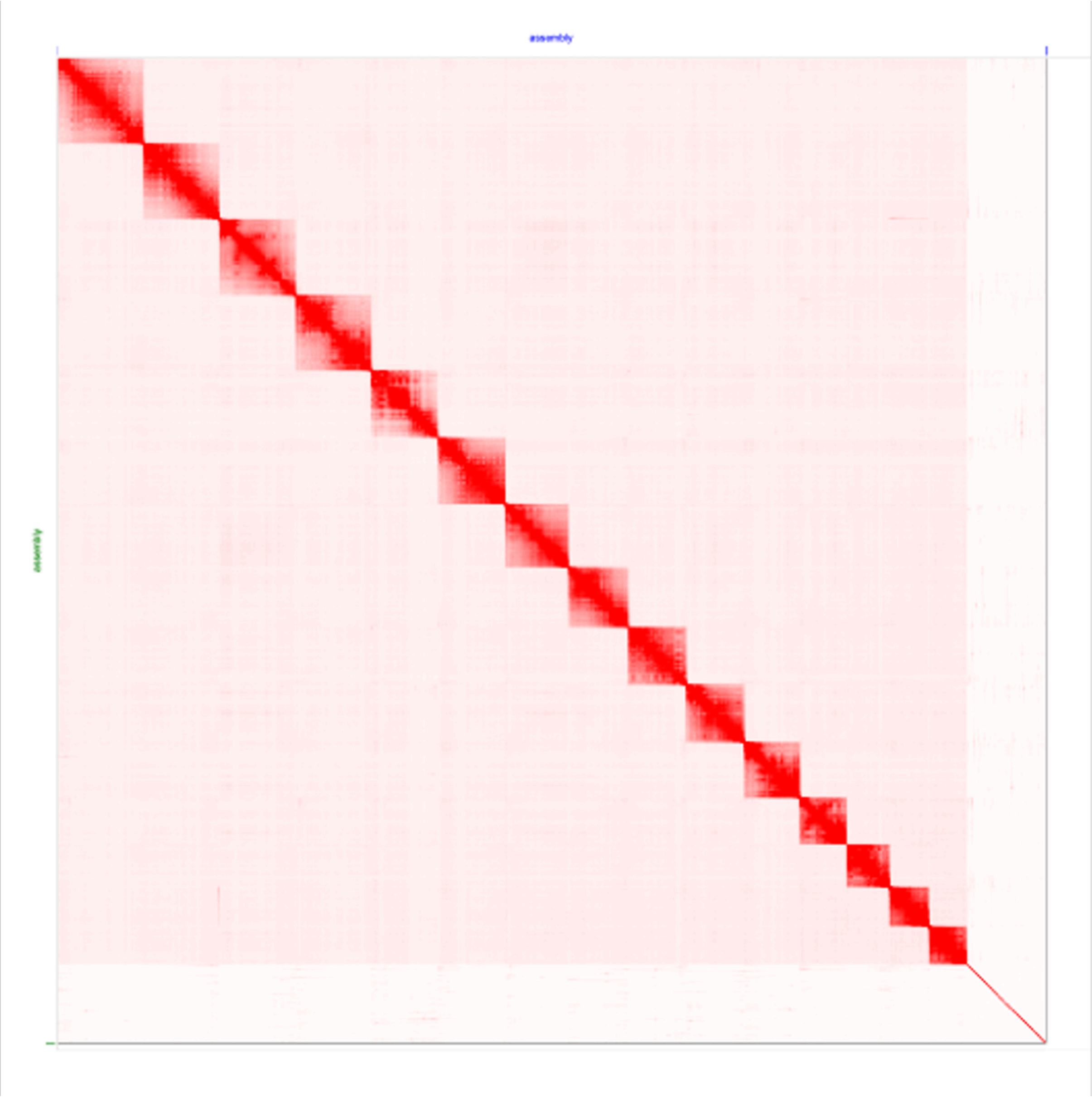

Figure S1. Hi-C contact map of the *S. africanus* assembly visualized using Juicebox. The intensity of the red signal denotes Hi-C contact density between assembly regions. Each high-contact block corresponds to a scaffolded chromosome.

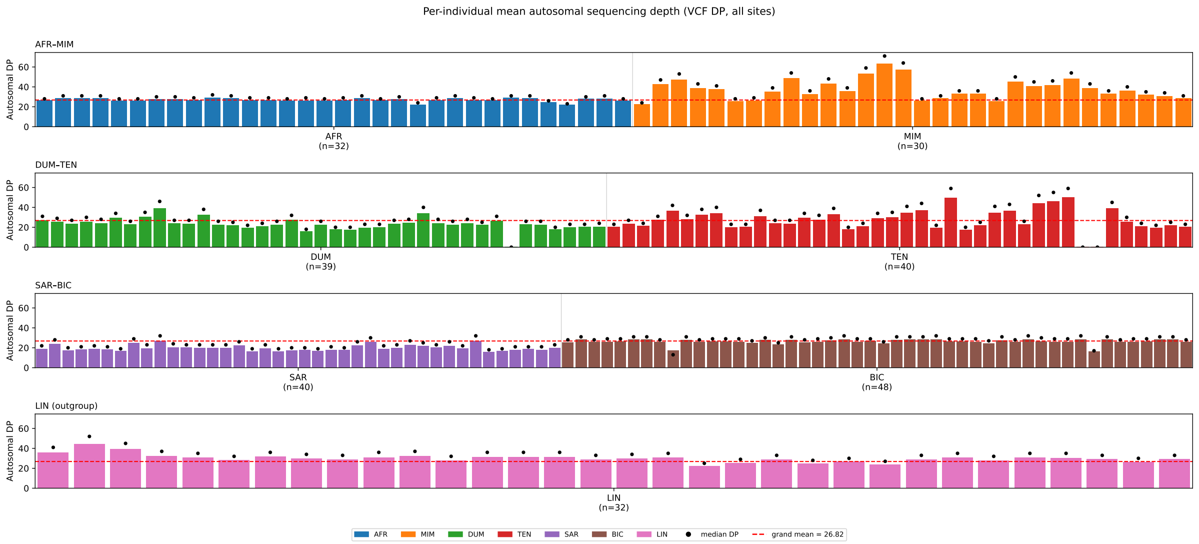

Figure S2. The sequence coverage across individuals per species based on the called genotype. Species in social-subsocial pairs are visualised in the same panel. Each bar and dot represents the average and median genotype depth per individual accordingly. The red dashed line denotes the mean genotype depth across all genotyped individuals.

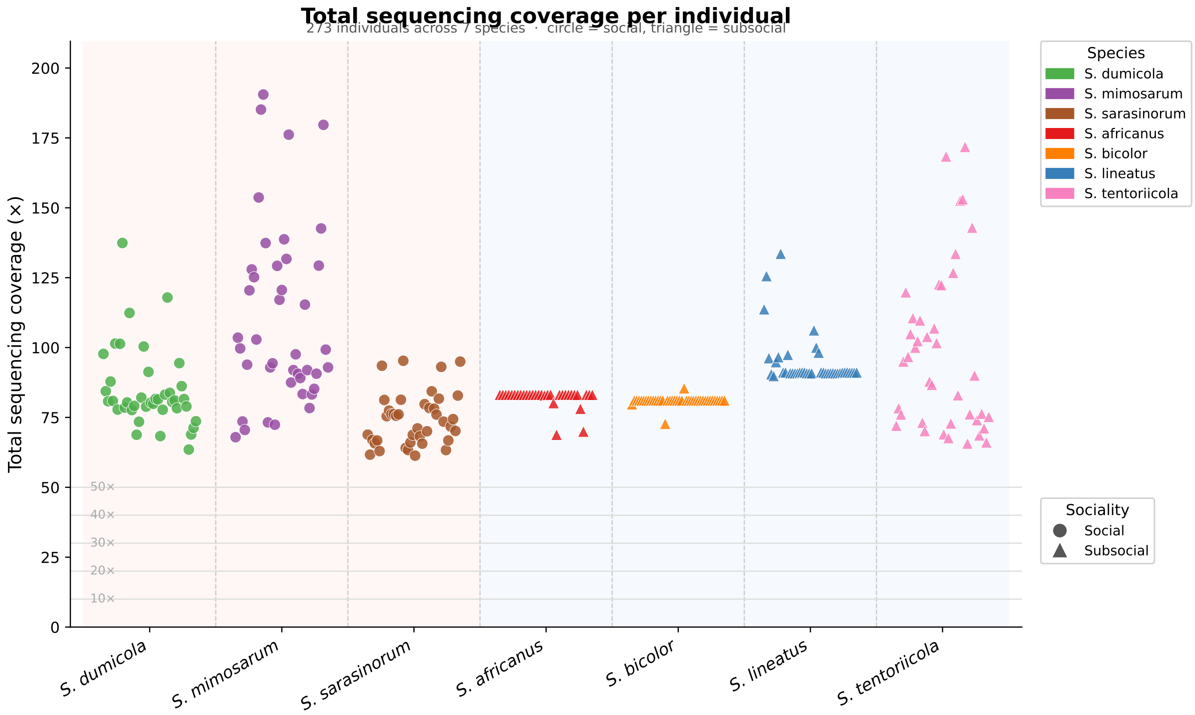

Figure S3. Raw whole-genome sequencing depth across all individuals. The figure summarizes total raw read depth per individual across species, families, and parent/offspring roles to show that sequencing depth was comparable across the dataset.

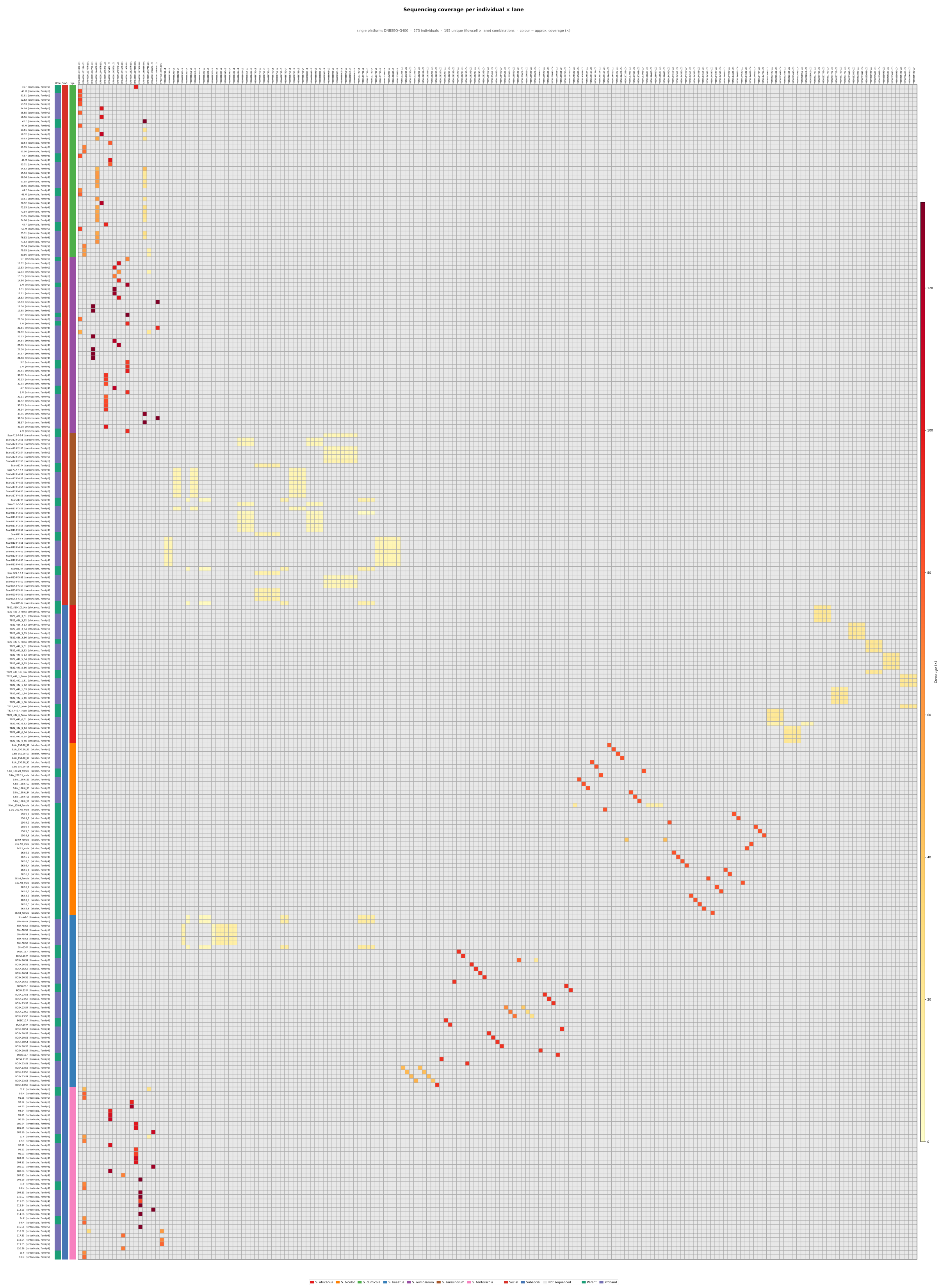

Figure S4. The heatmap of sequencing flowcell and lane assignment across individuals. The figure visualizes per-individual species, family, parent/offspring sample, flowcell IDs, lanes, and total coverage to show that sequencing was not organized into large biological-status or role-specific batches.

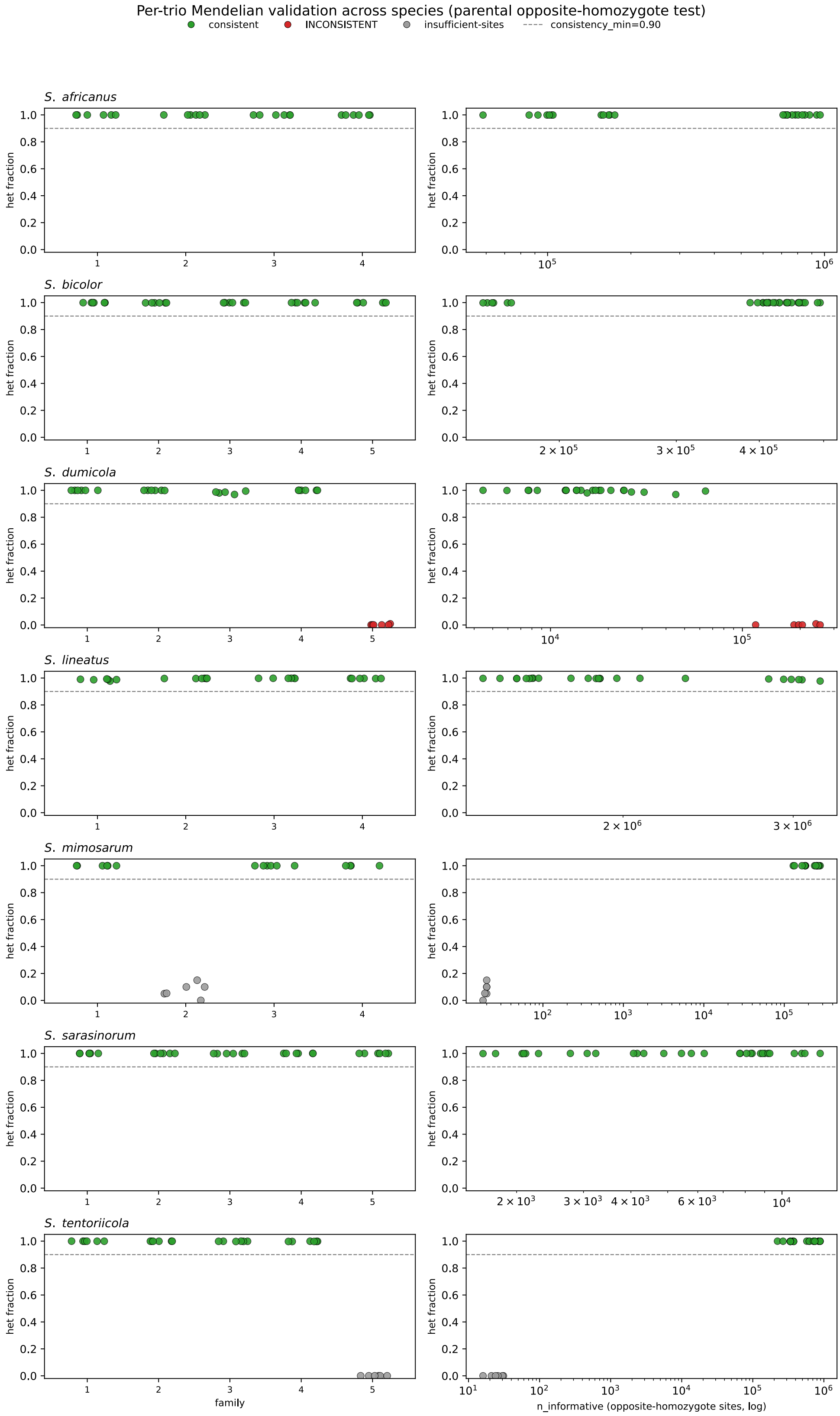

777 Figure S5. Kinship verification across sequenced trios. Mendelian consistency scores were calculated
778 from autosomal biallelic SNPs where the two assigned parents were homozygous for opposite alleles.
779 Correctly assigned trios are expected to show near-complete offspring heterozygosity at these
780 informative sites.

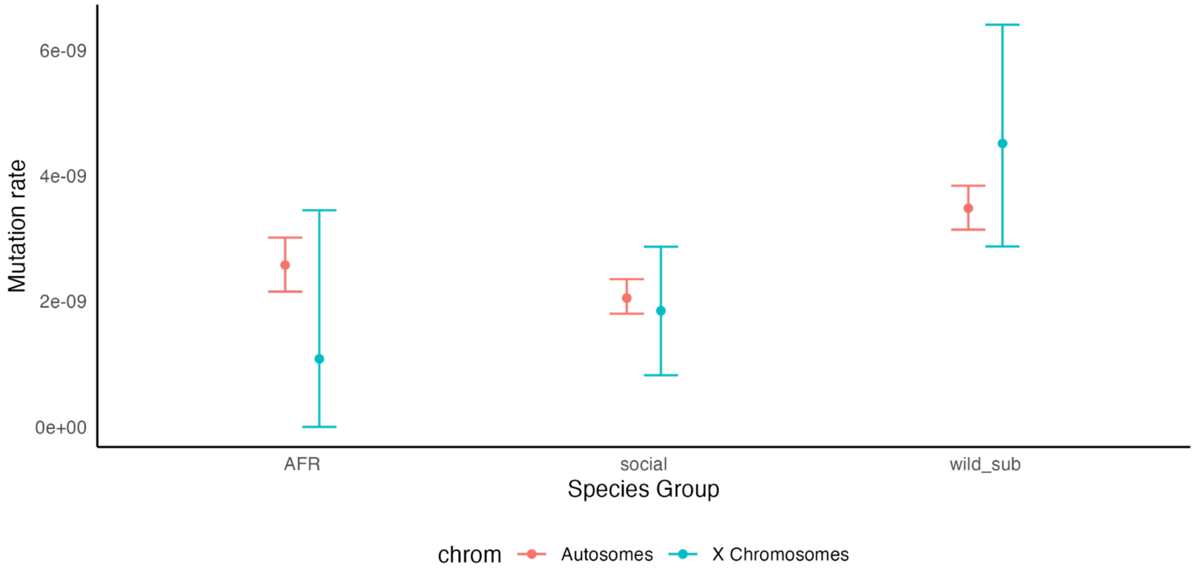

Figure S6. Autosomal and X chromosome mutation-rate estimates under the final minimum-depth filter. AFR denotes *S. africanus*, social denotes *S. dumicola*, *S. mimosarum*, and *S. sarasinorum*, and wild_sub denotes *S. lineatus*, *S. tentoriicola,* and *S. bicolor* with hypermutated families excluded.

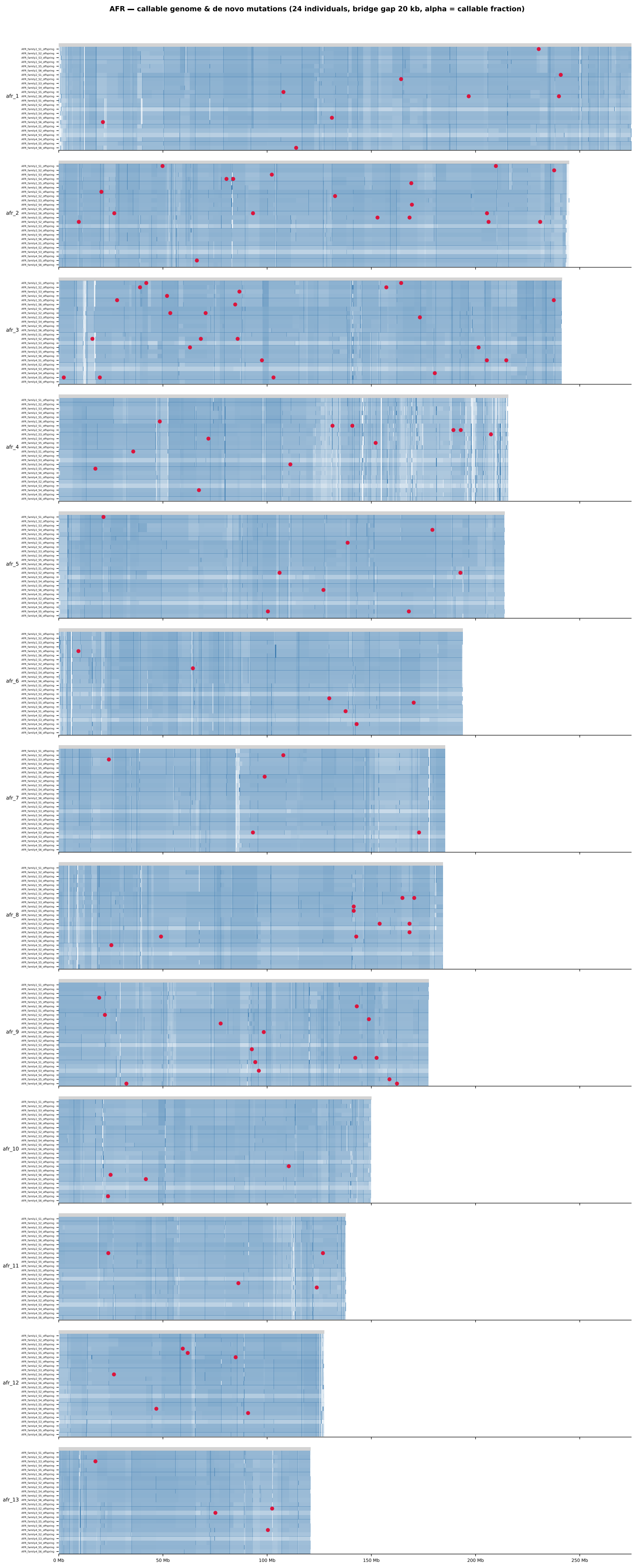

Figure S7. Genomic distribution of detected autosomal DNMs in *S. africanus*.

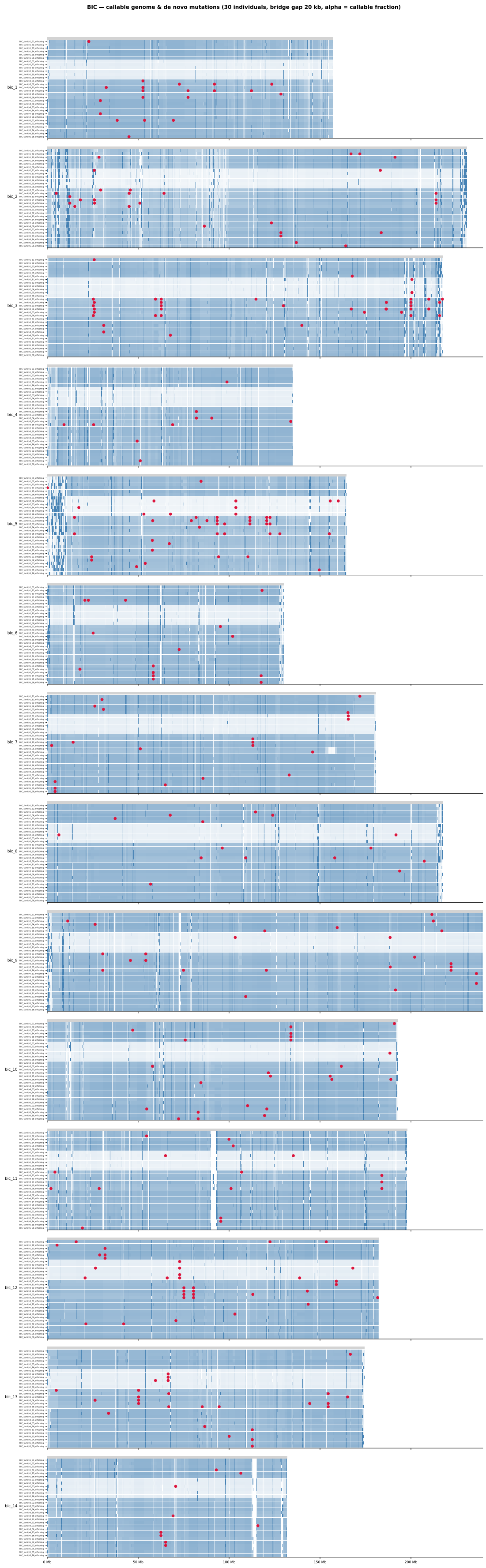

Figure S8. Genomic distribution of detected autosomal DNMs in *S. bicolor*.

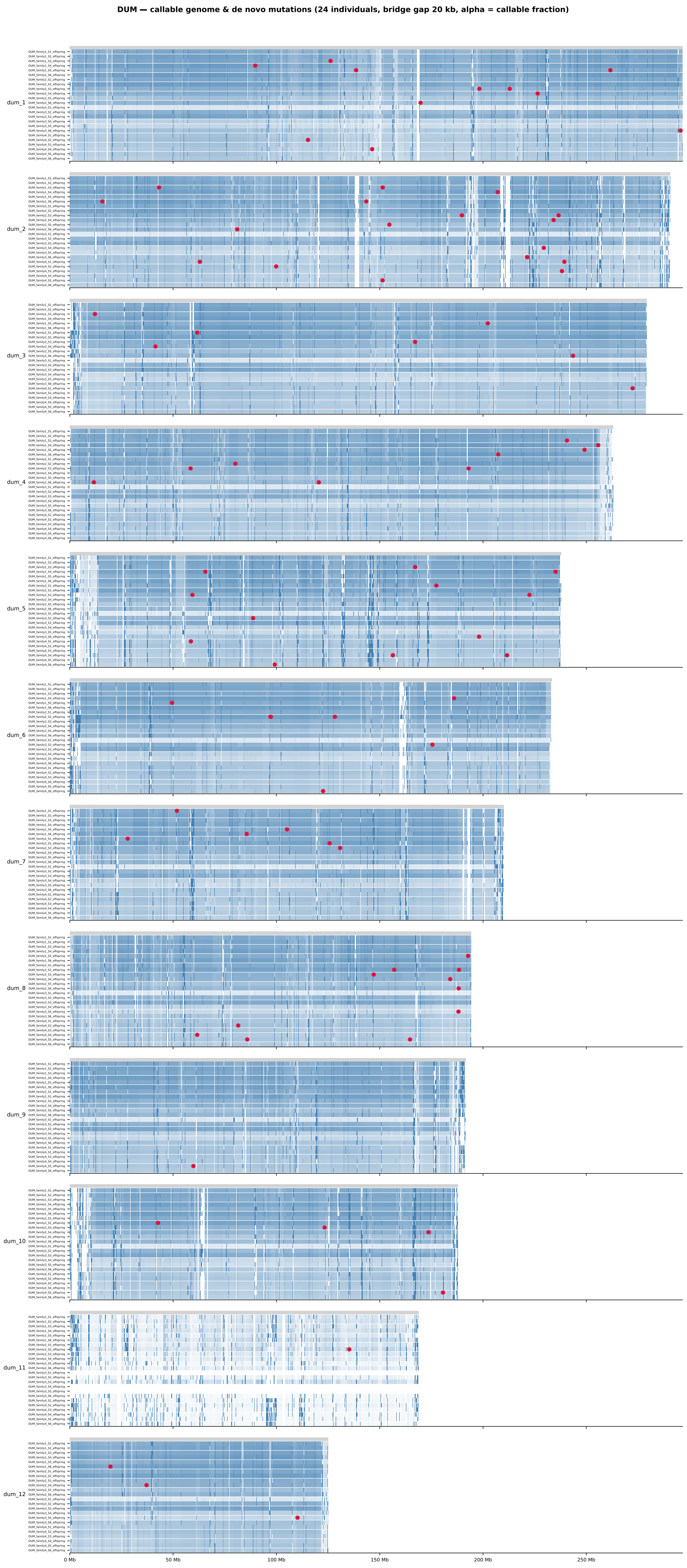

Figure S9. Genomic distribution of detected autosomal DNMs in *S. dumicola*.

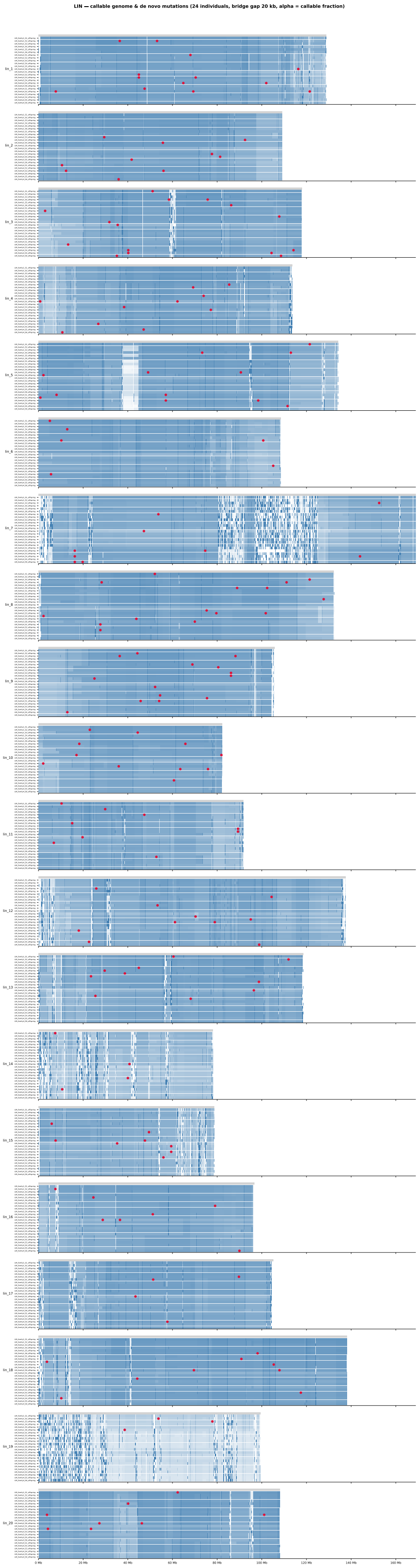

Figure S10. Genomic distribution of detected autosomal DNMs in *S. lineatus*.

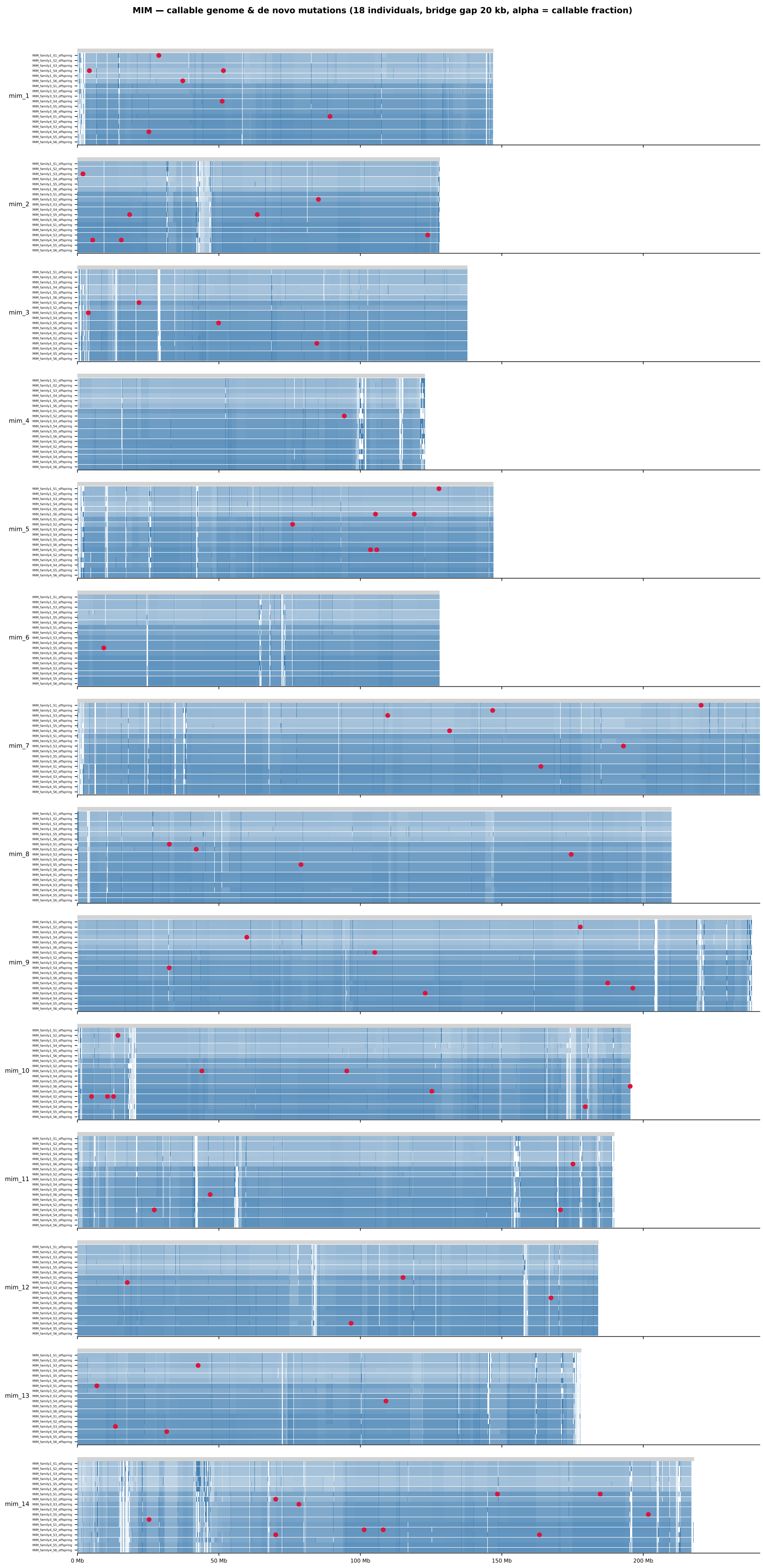

788 Figure S11. Genomic distribution of detected autosomal DNMs in *S. mimosarum*.

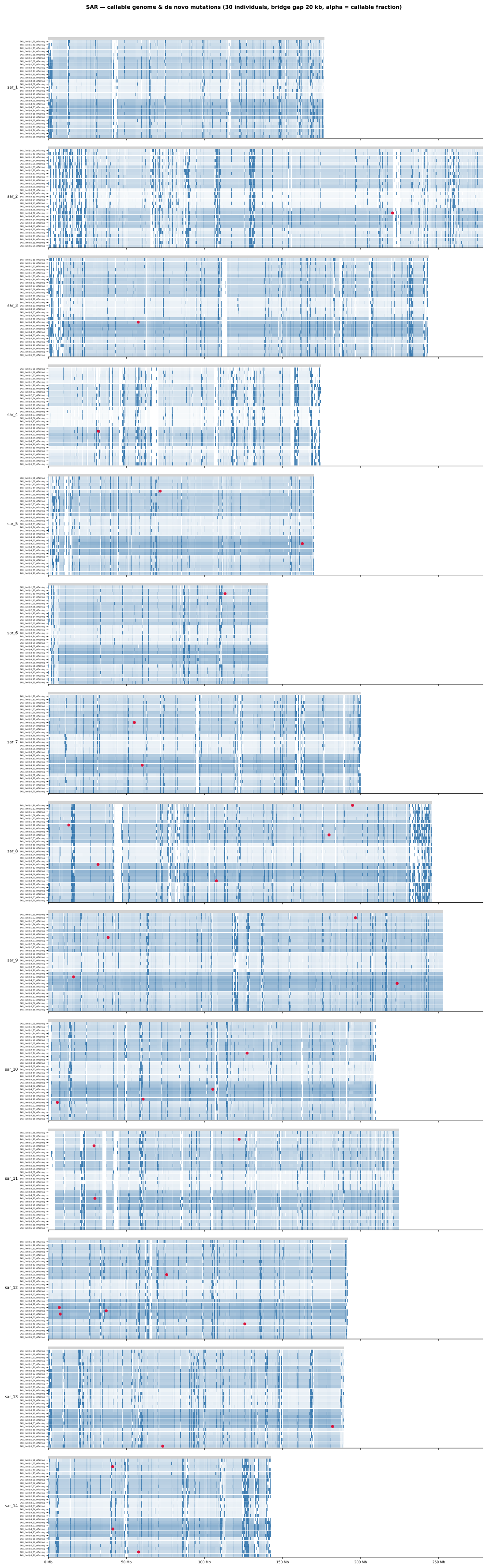

789 Figure S12. Genomic distribution of detected autosomal DNMs in *S. sarasinorum*.

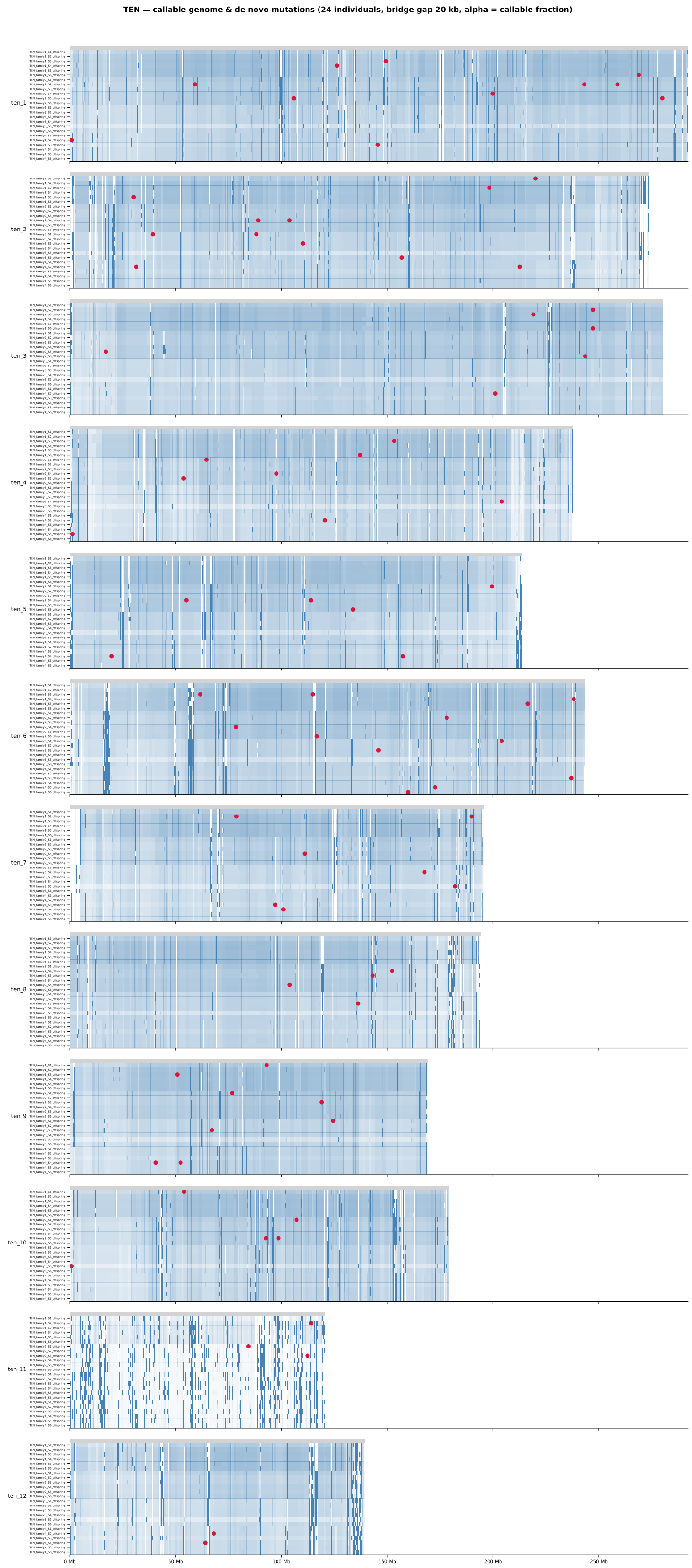

Figure S13. Genomic distribution of detected autosomal DNMs in *S. tentoriicola*. For Figures S3-S9, each panel represents a chromosome with DNMs marked as red points. Each row within a panel corresponds to a sampled trio. The blue gradient indicates local callability, with darker blue denoting higher callability and white denoting non-callable regions.

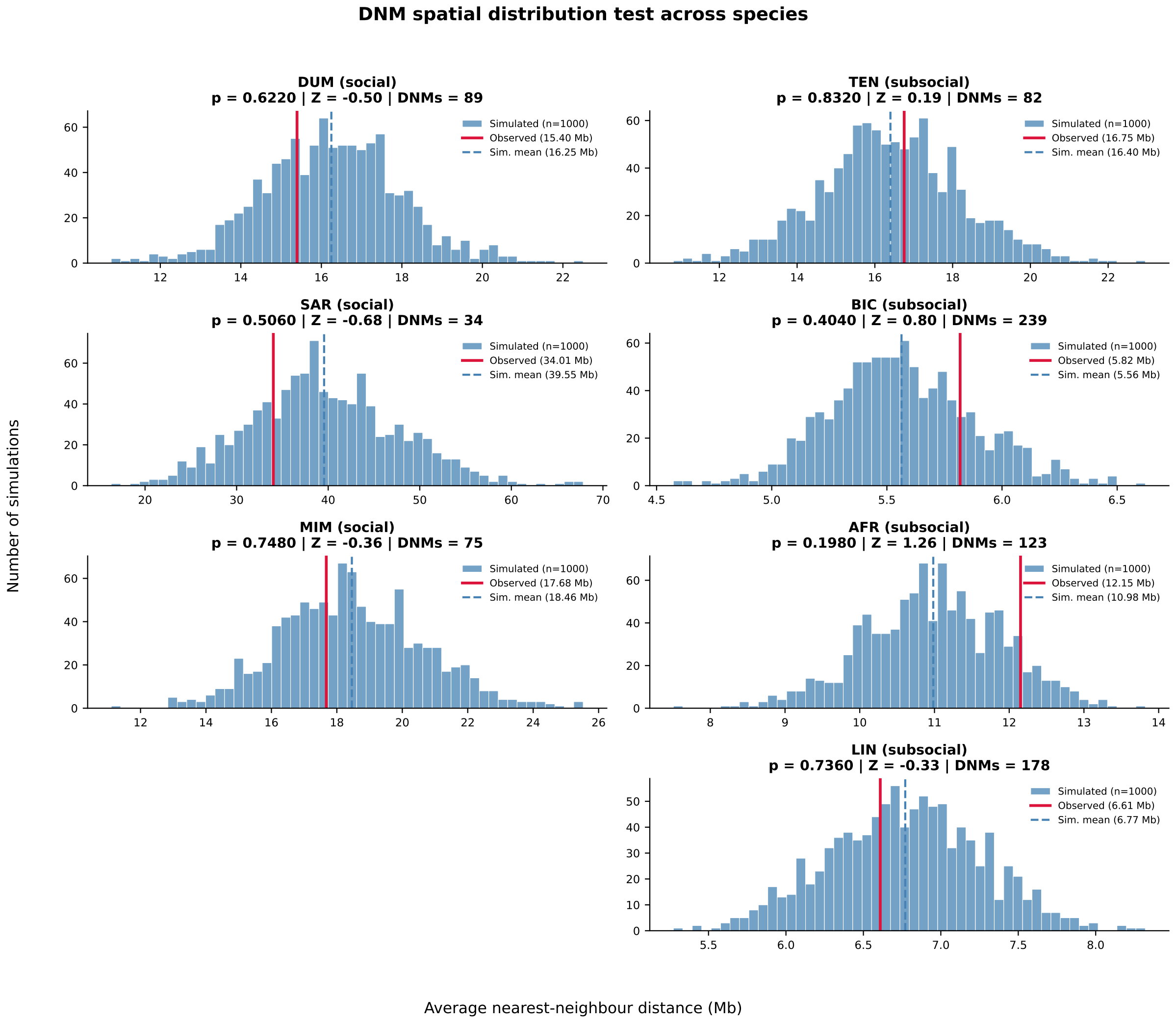

Figure S14. Testing observed nearest-neighbor distances among detected DNMs against a bootstrapped null distribution under a uniform mutation-rate model across callable regions.

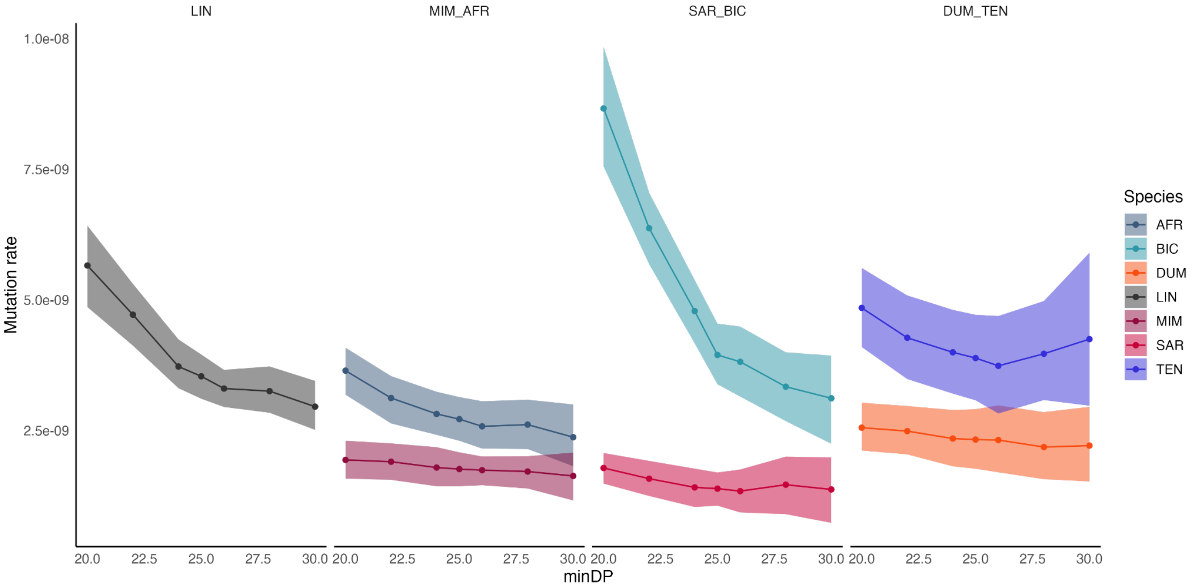

Figure S15. Bootstrapped mutation-rate estimates per species under different minimum-depth thresholds for callable-site filtering. The outgroup species S. lineatus and the three social-subsocial species pairs are shown in separate facets.

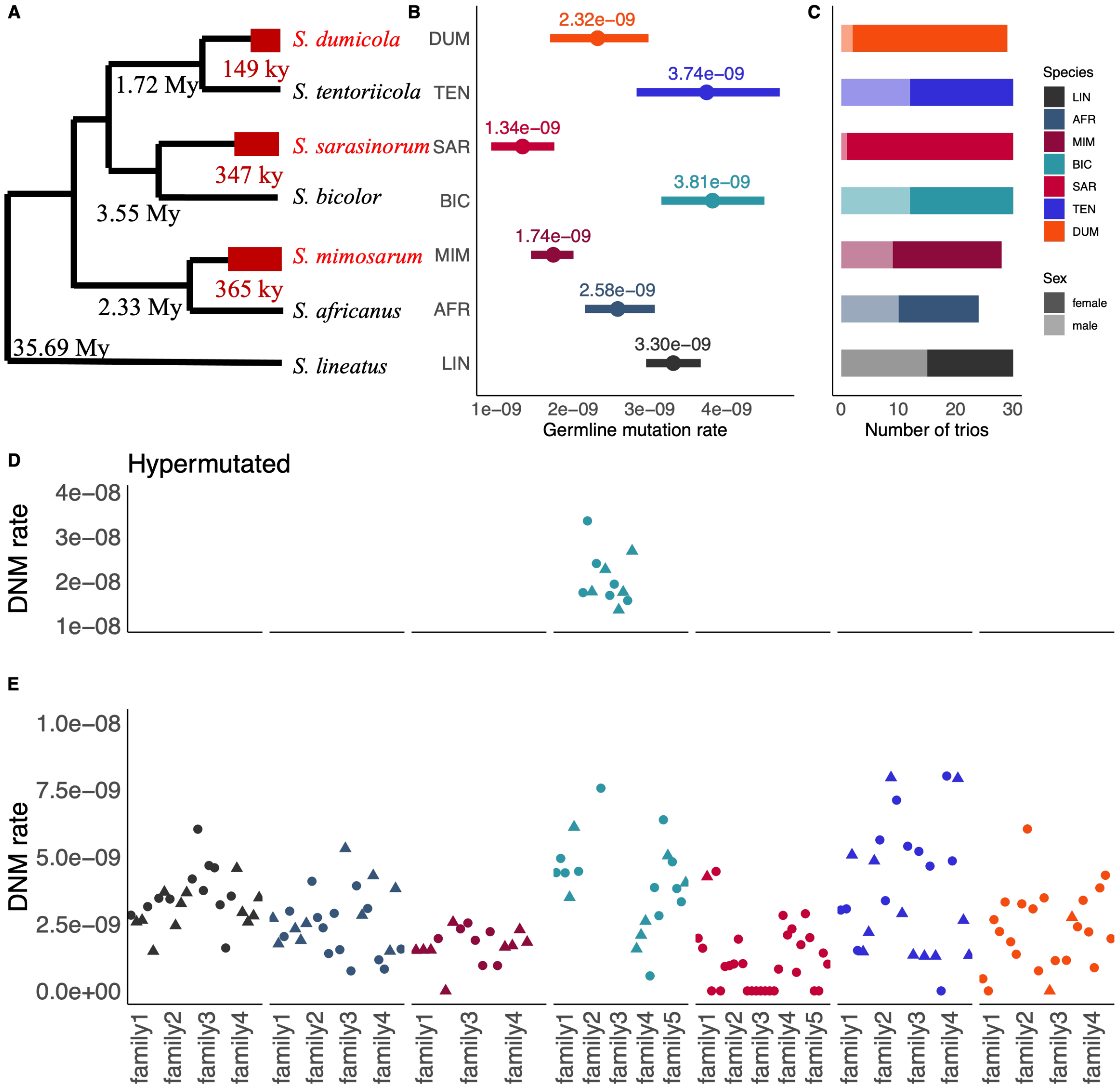

799 Figure S16. Mutation rate observations including hypermutated families. The layout resembles main
800 Figure 1 with hypermutated trios of *S. bicolor* visualised in the panel D.

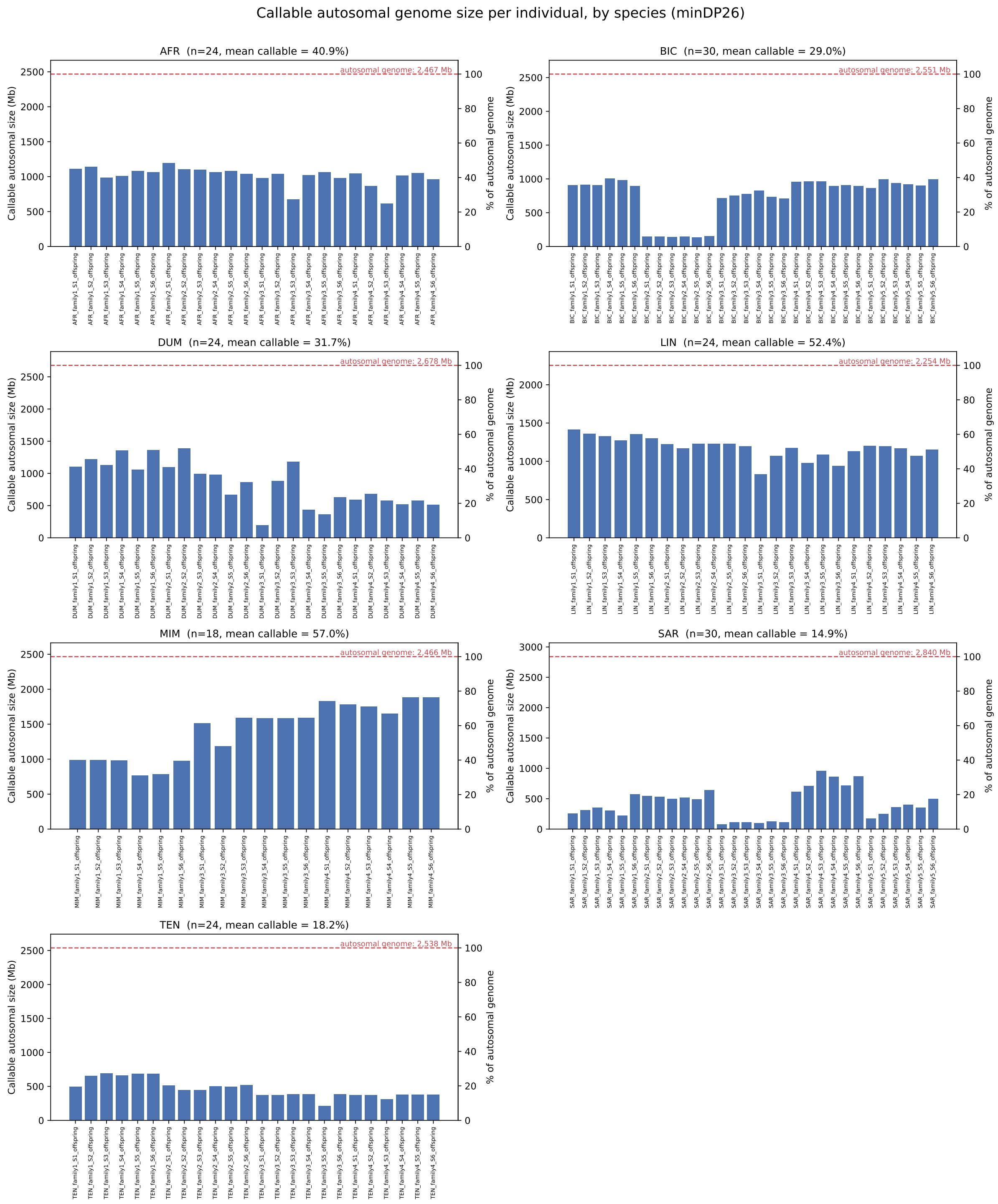

Figure S17. Callable genome size per trio based on autosomes. The total autosomal callable genome size is shown on the y-axis on the left side, and the callable genome fraction is shown on the y-axis on the right side.

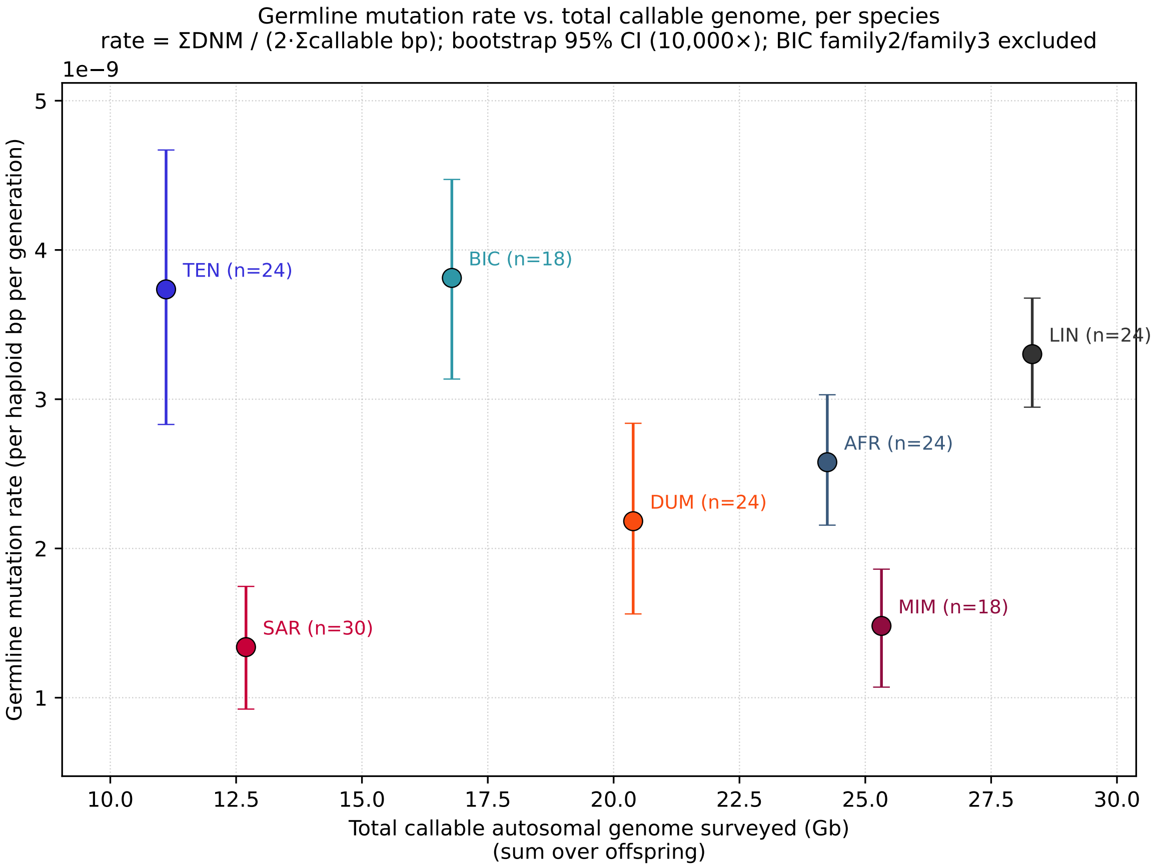

Figure S18. The total callable autosomal genome and the corresponding germline mutation rate estimate per species.

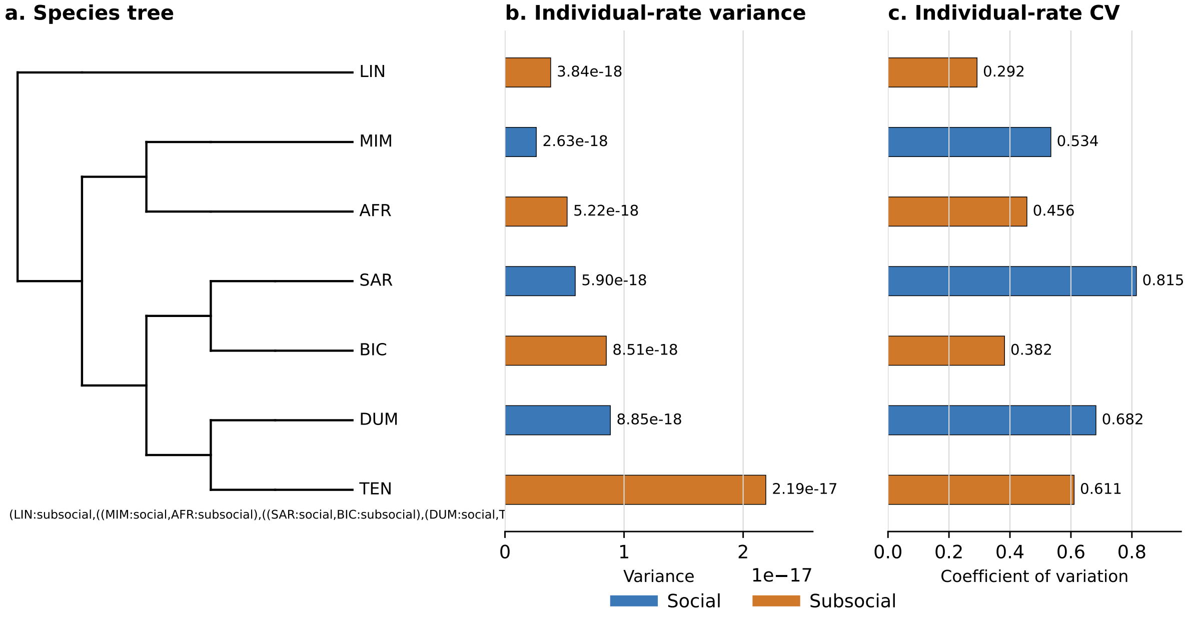

806 Figure S19. Variance and coefficient of variation of individual mutation-rate estimates per species.

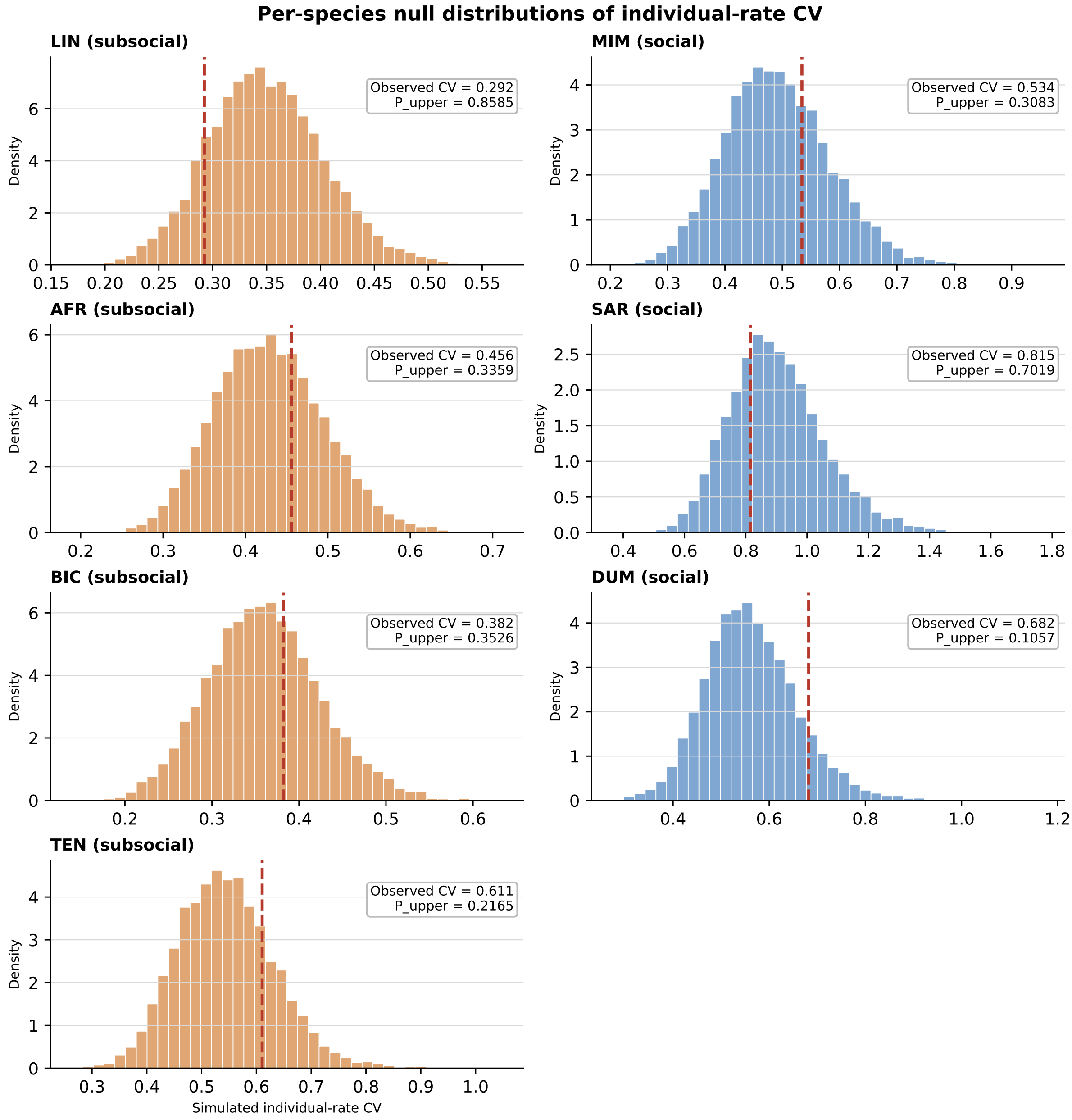

Figure S20. Observed coefficient of variation compared with simulated null distributions assuming a constant species-specific mutation rate.

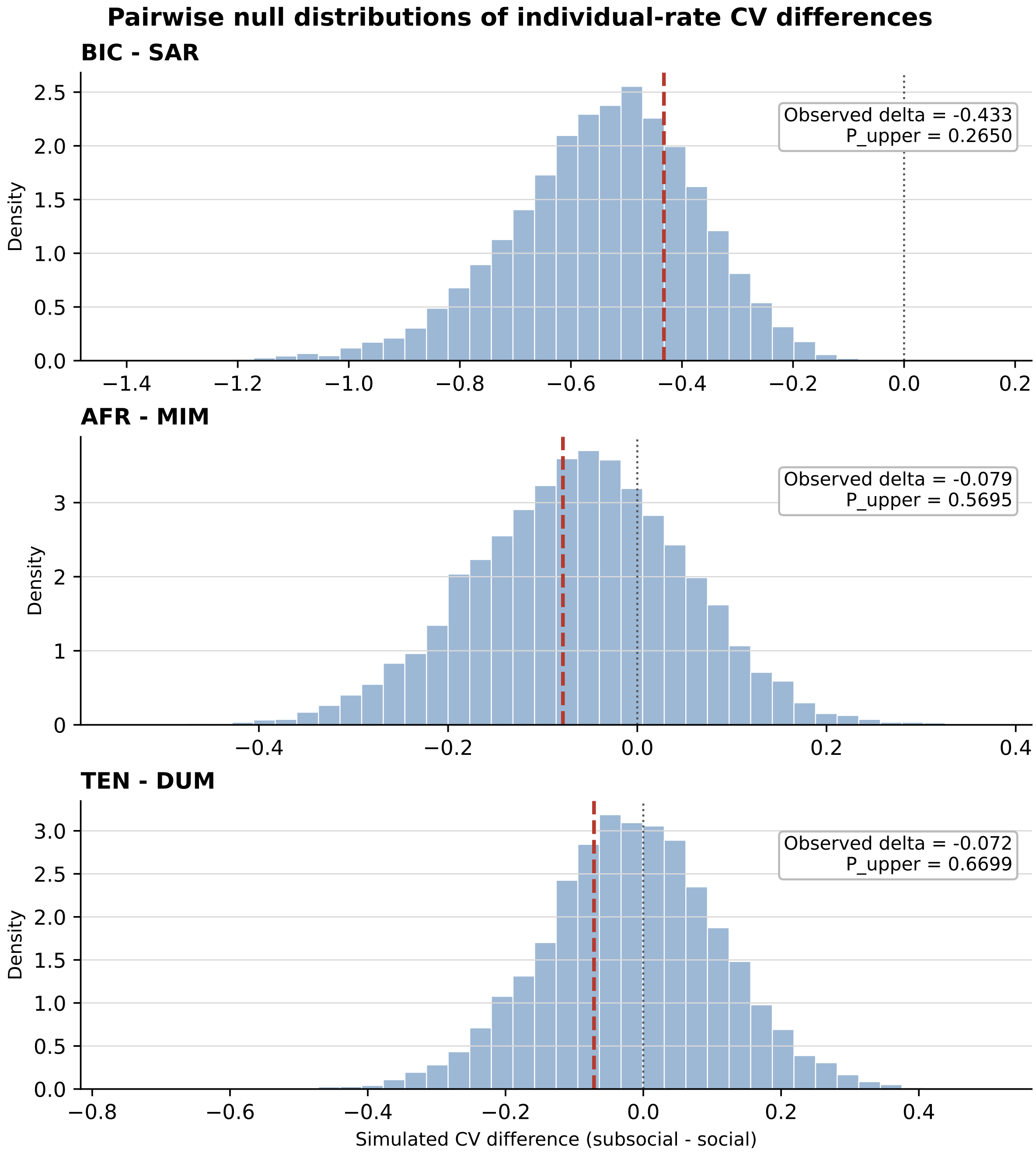

Figure S21. Observed pairwise differences in the coefficient of variation compared with simulated null distributions for each social-subsocial species pair.

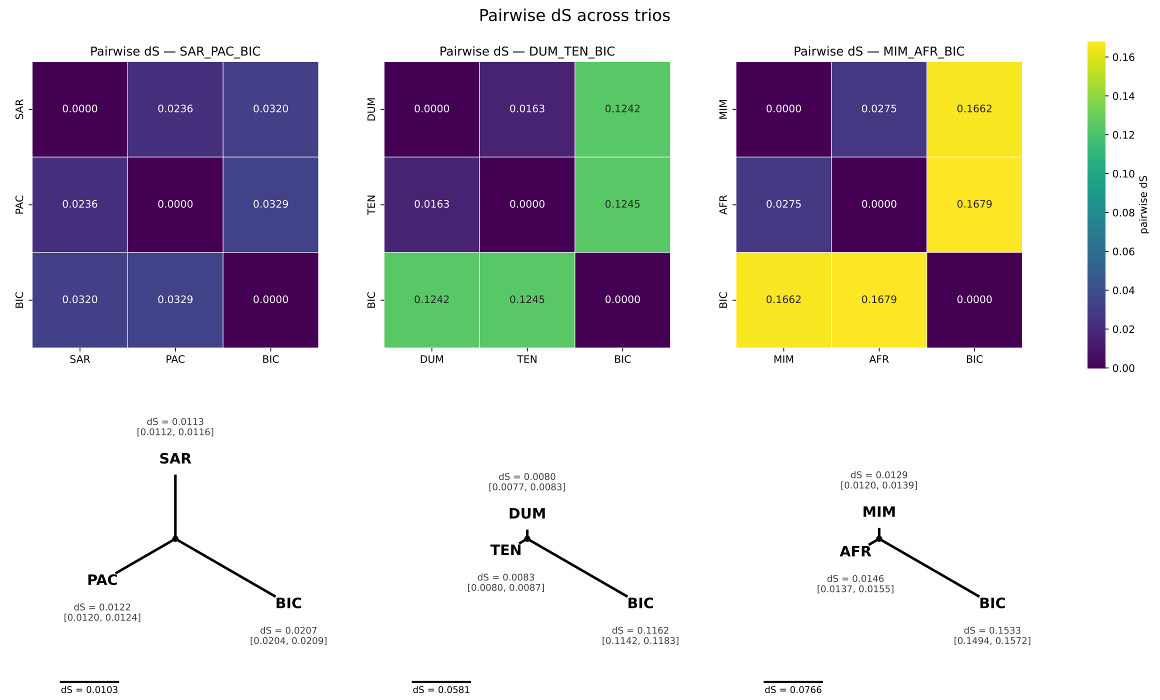

Figure S22. Pairwise $d_s$ estimates and derived lineage branch lengths for social-subsocial species pairs using short-read DNA aligned to the S. bicolor reference genome.

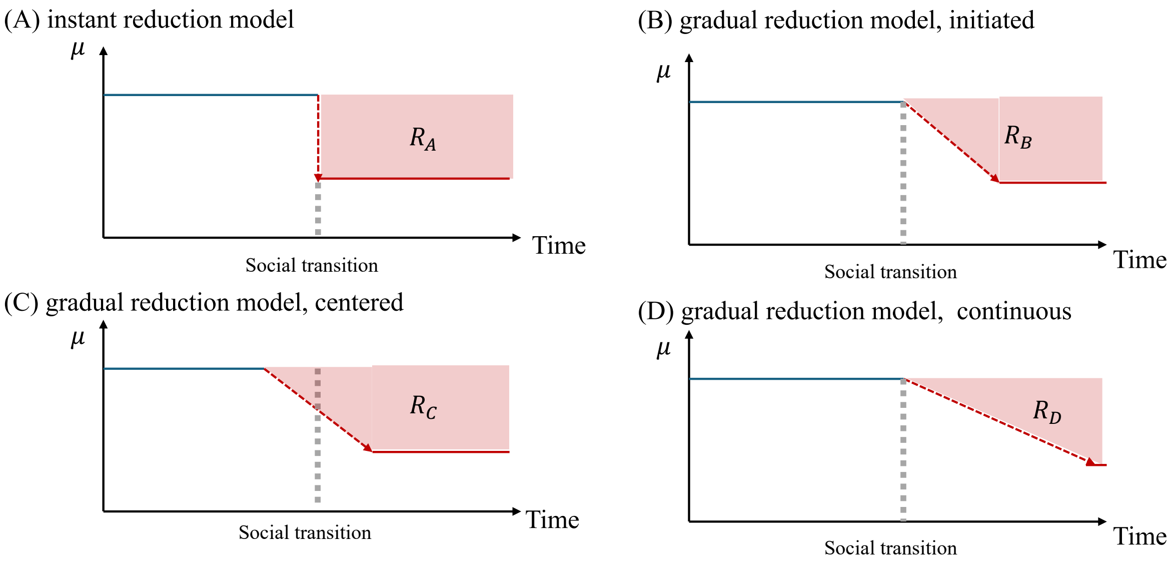

$$R_A = R_C > R_B > R_D = \frac{R_A}{2}$$

Figure S23. Gradual mutation-rate reduction model. The schematic illustrates expected reductions in accumulated $d_S$ under instantaneous and gradual reductions in mutation rate around the inferred transition to sociality. Gradual reductions produce an expected $d_S$ reduction between the instantaneous-shift expectation and approximately one half of that expectation, depending on whether the rate change begins at, or is centered on, the social transition.

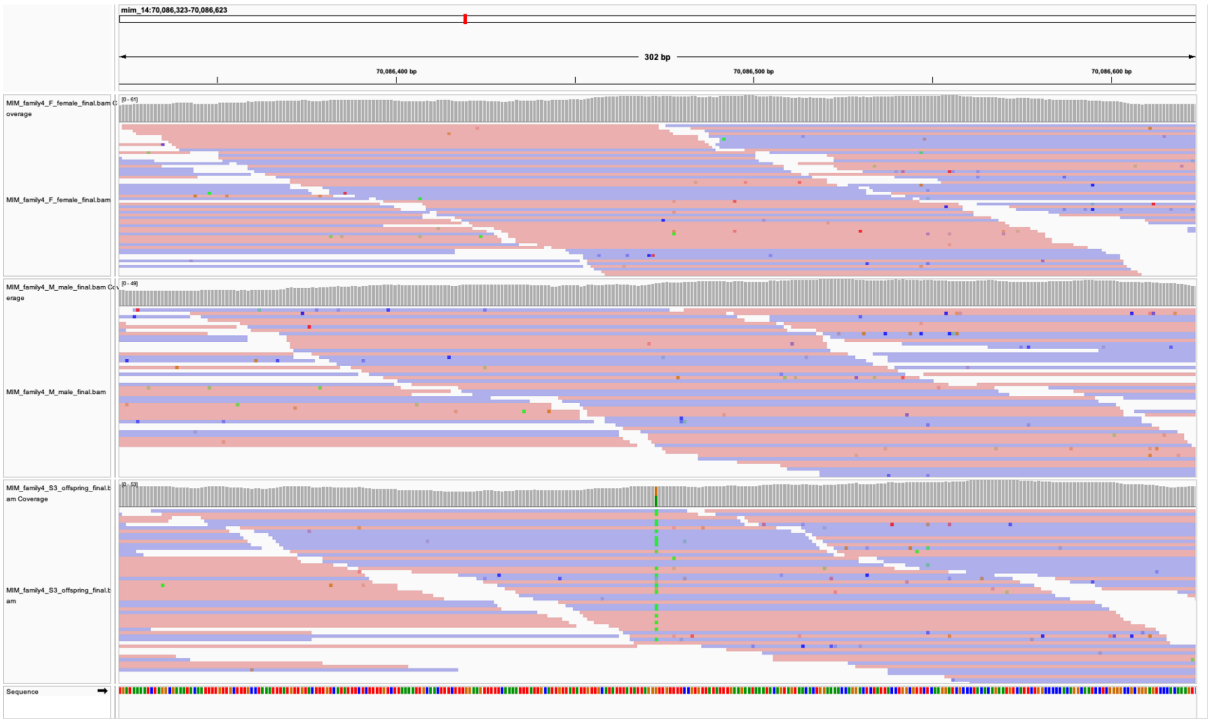

Figure S24. IGV inspection of the shared DNM between offspring from different *S. mimosarum* families: MIM_family4_S3_offspring, mim_14:70086473.

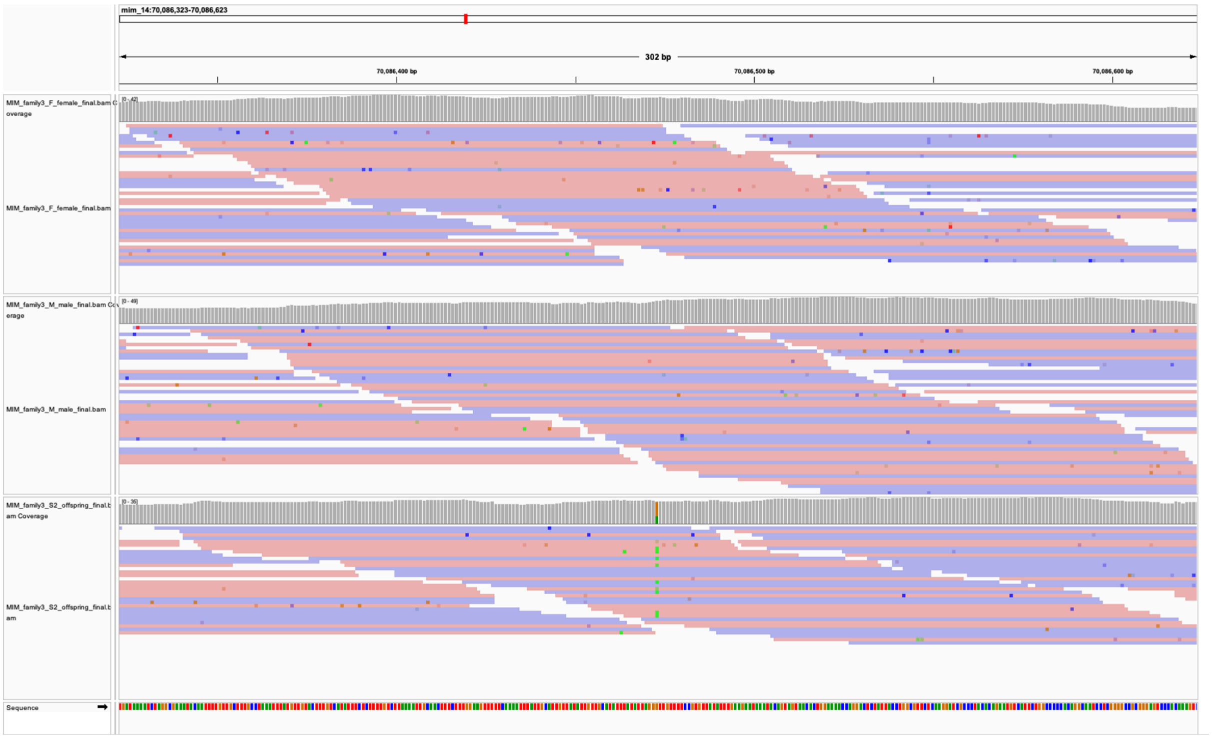

Figure S25. IGV inspection of the shared DNM between offspring from different *S. mimosarum* families: MIM_family3_S2_offspring, mim_14:70086473.

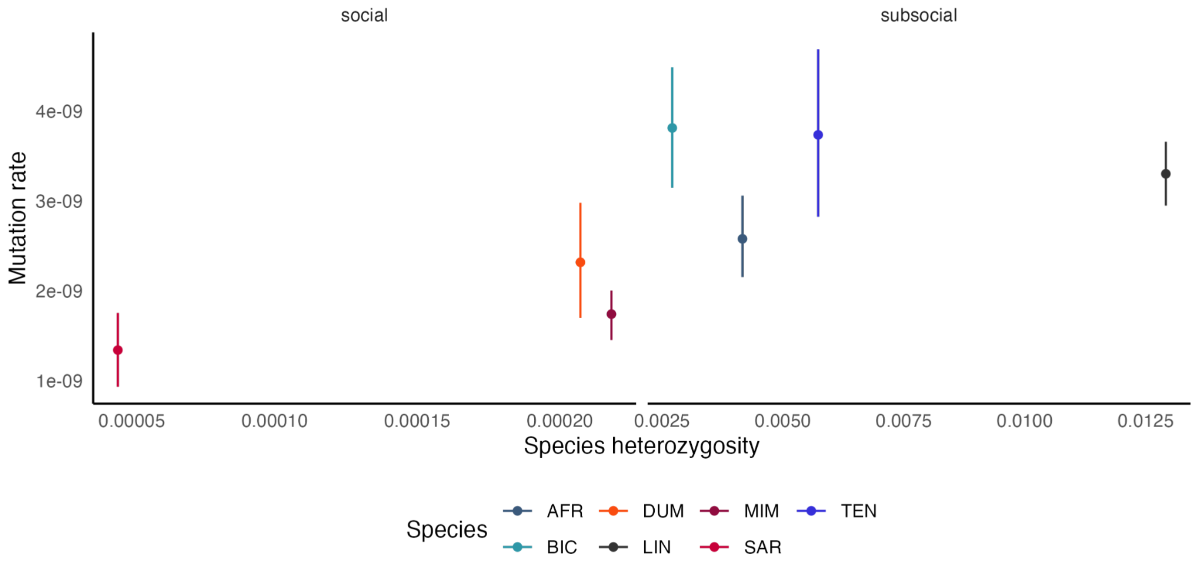

Figure S26. Relationship between species heterozygosity and estimated mutation rate. The x-axis shows mean heterozygosity per species based on individual heterozygosity estimates, and the y-axis shows estimated mutation rate.

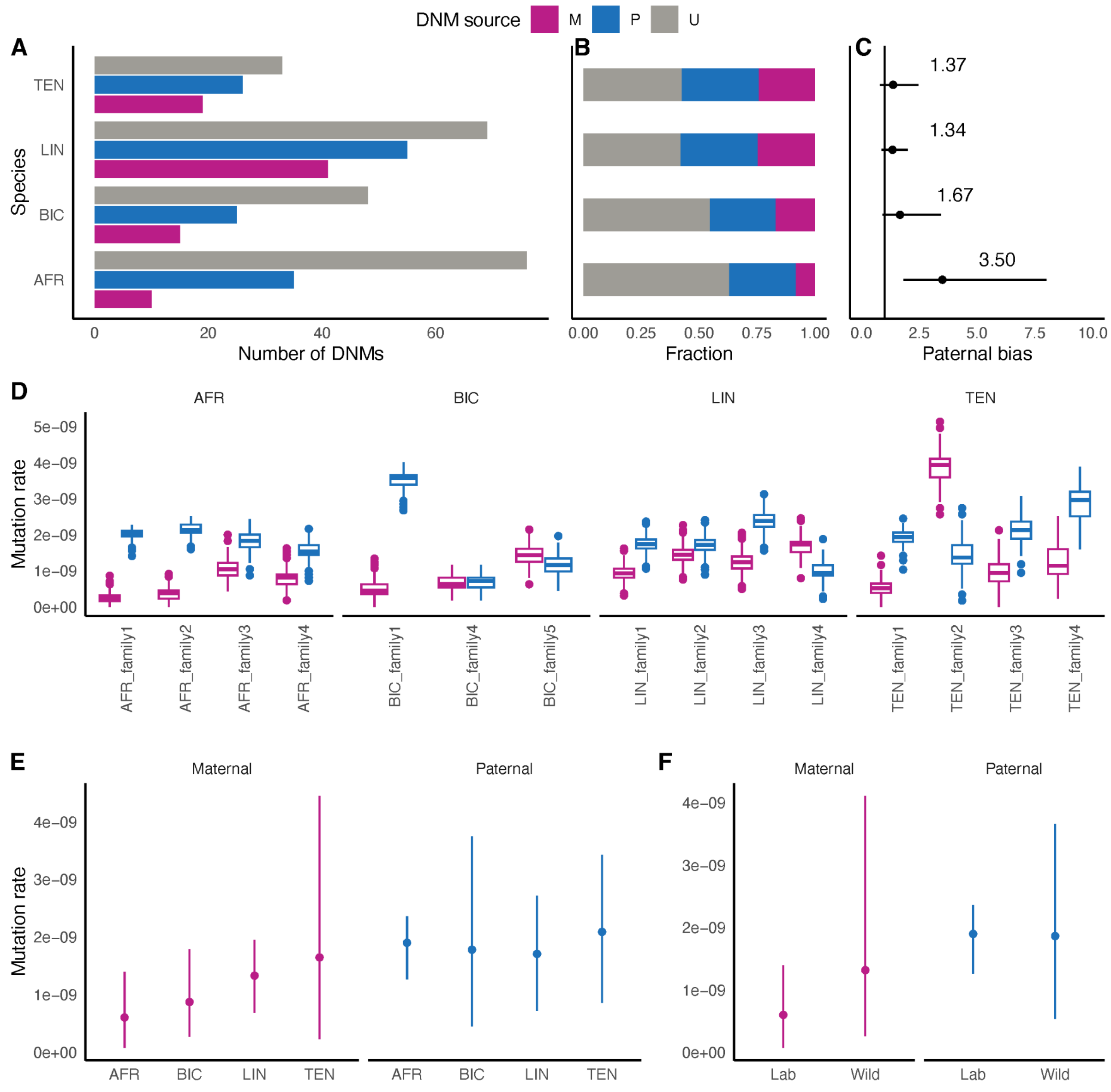

Figure S27. Paternal mutation bias in subsocial spider species. (A) Number of DNMs with phased paternal or maternal origin and unphased DNMs for all subsocial species. (B) Fraction of phased DNMs per subsocial species. (C) Paternal bias estimated for each subsocial species based on phased DNMs. The vertical line denotes no paternal bias. (D) Bootstrapped paternal and maternal mutation rates per family. (E) Paternal and maternal germline mutation rates per species. (F) Paternal and maternal germline mutation rates for lab-reared *S. africanus* and wild-caught subsocial species.

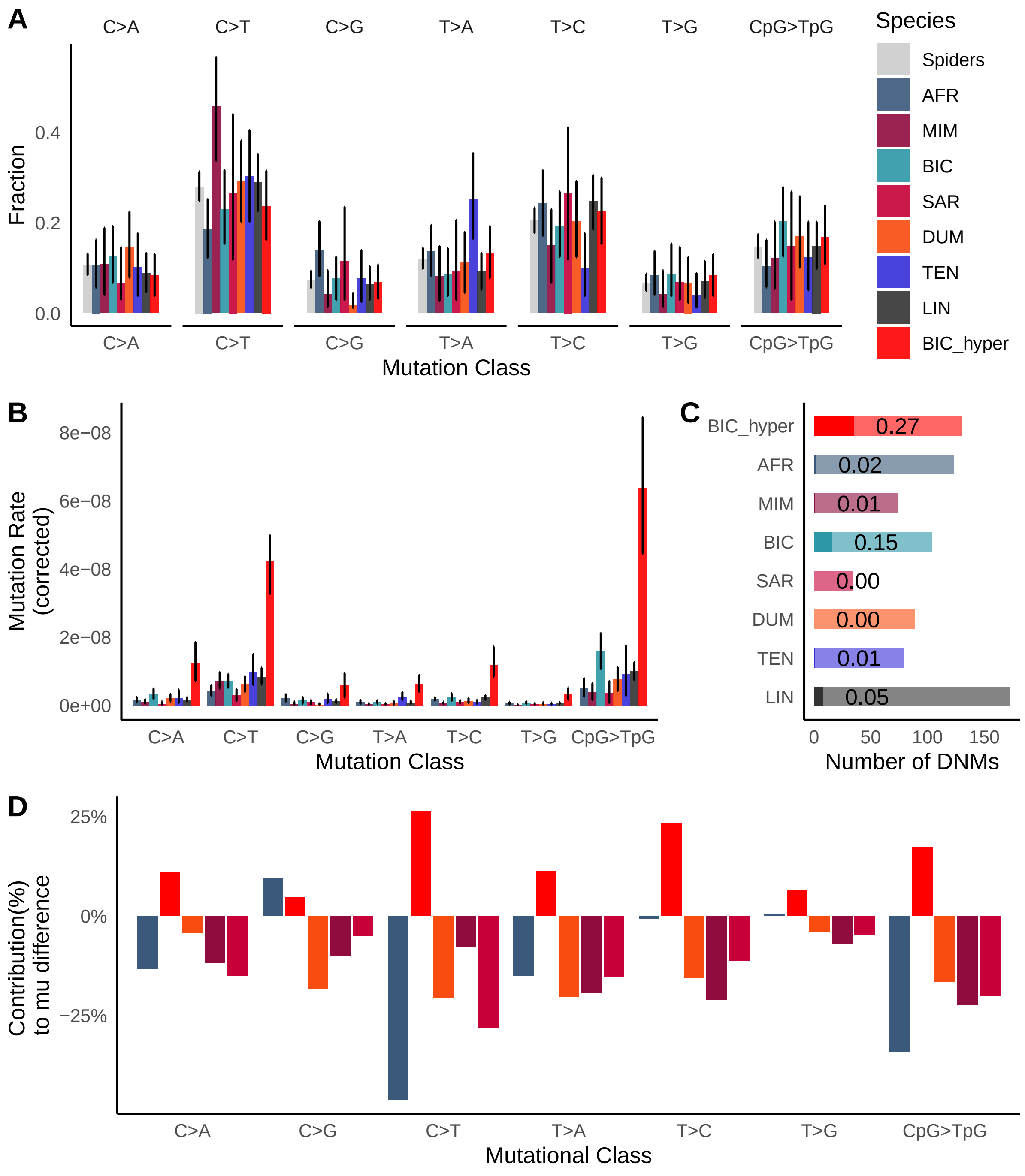

Figure S28. Mutational spectra comparison including hypermutated families. The layout resembles the main Figure 4 with data from hypermutated individuals being listed as a new separate category.

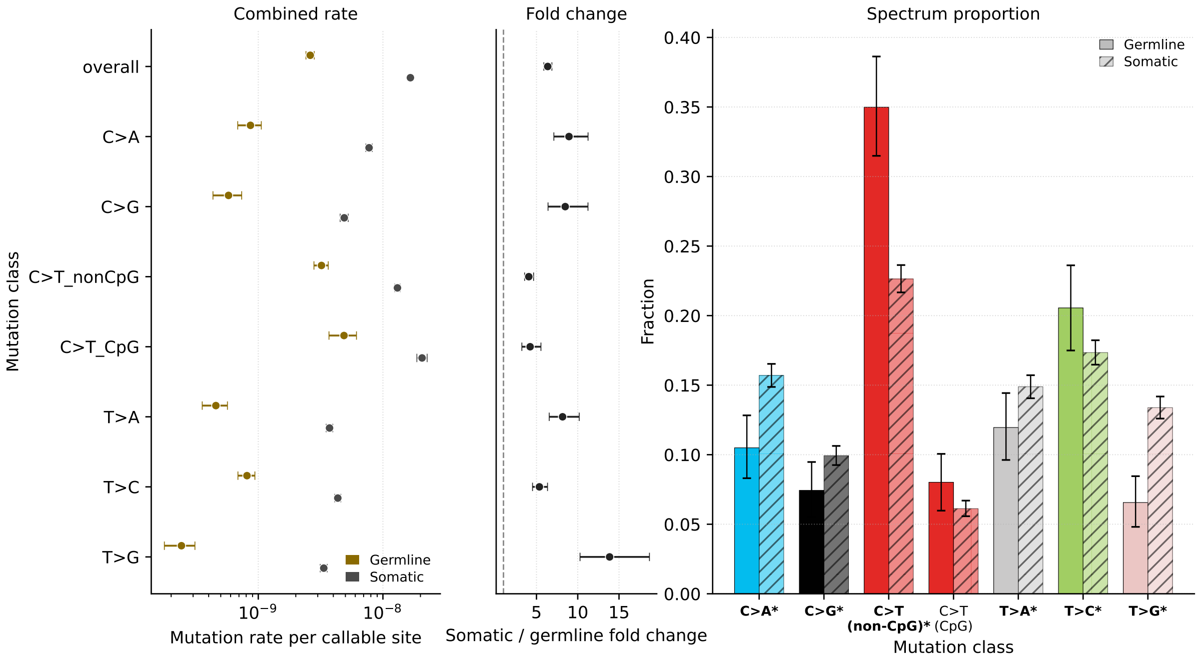

Figure S29. The comparison between somatic and germline mutation rate and mutational spectra. (a) The mutation rate per callable site in both germline mutations and somatic mutations spread out by the nucleotide context. (b) The mutation rate fold-change difference between the somatic mutation rate and the germline mutation rate. (c) The comparison between the mutational spectra of the germline mutations and the somatic mutations, mutation categories with significant fisher's exact test results are highlighted in the bold font.

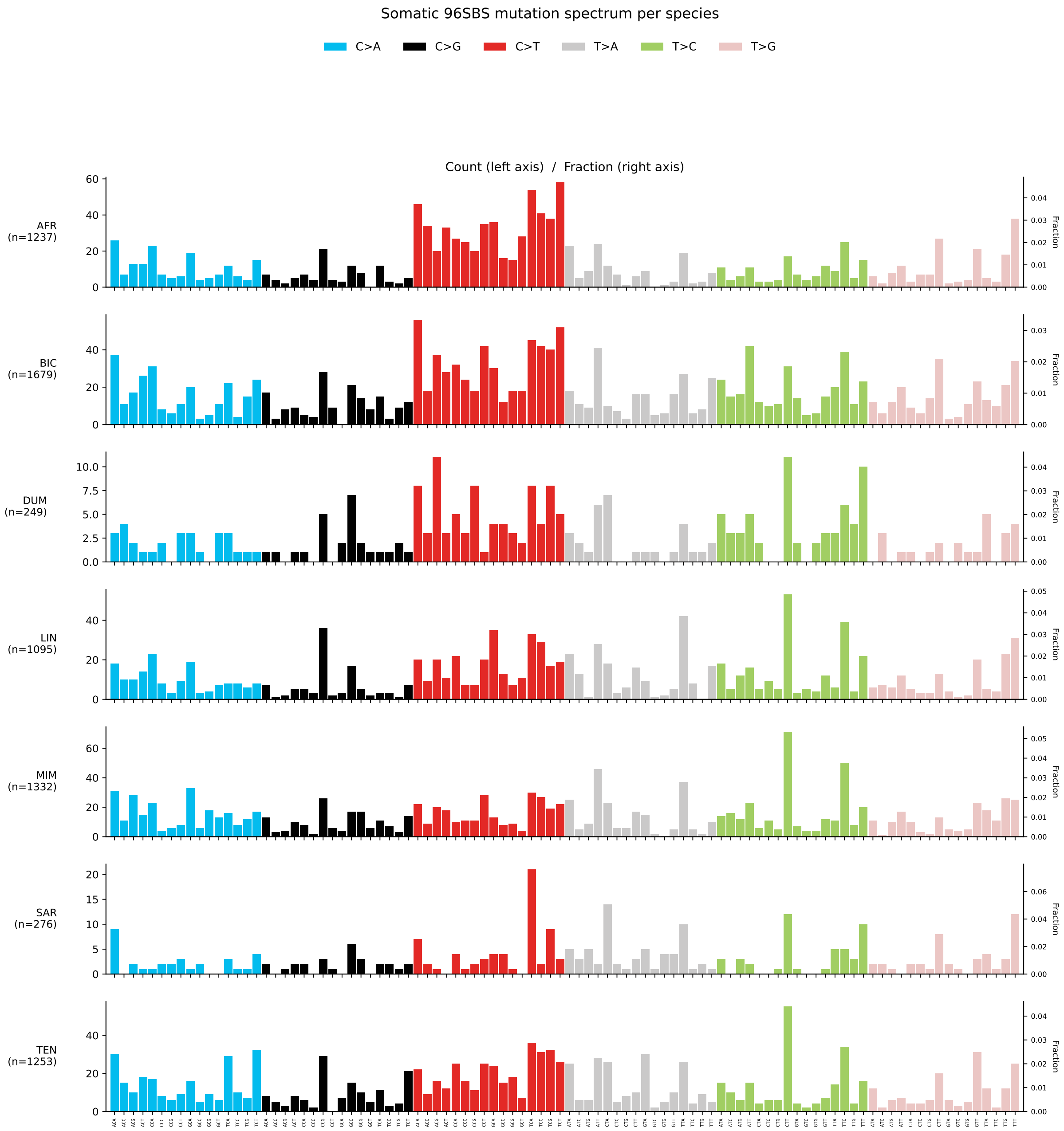

839 Figure S30. The somatic mutation spectra with trinucleotide context across all *Stegodyphus* species.

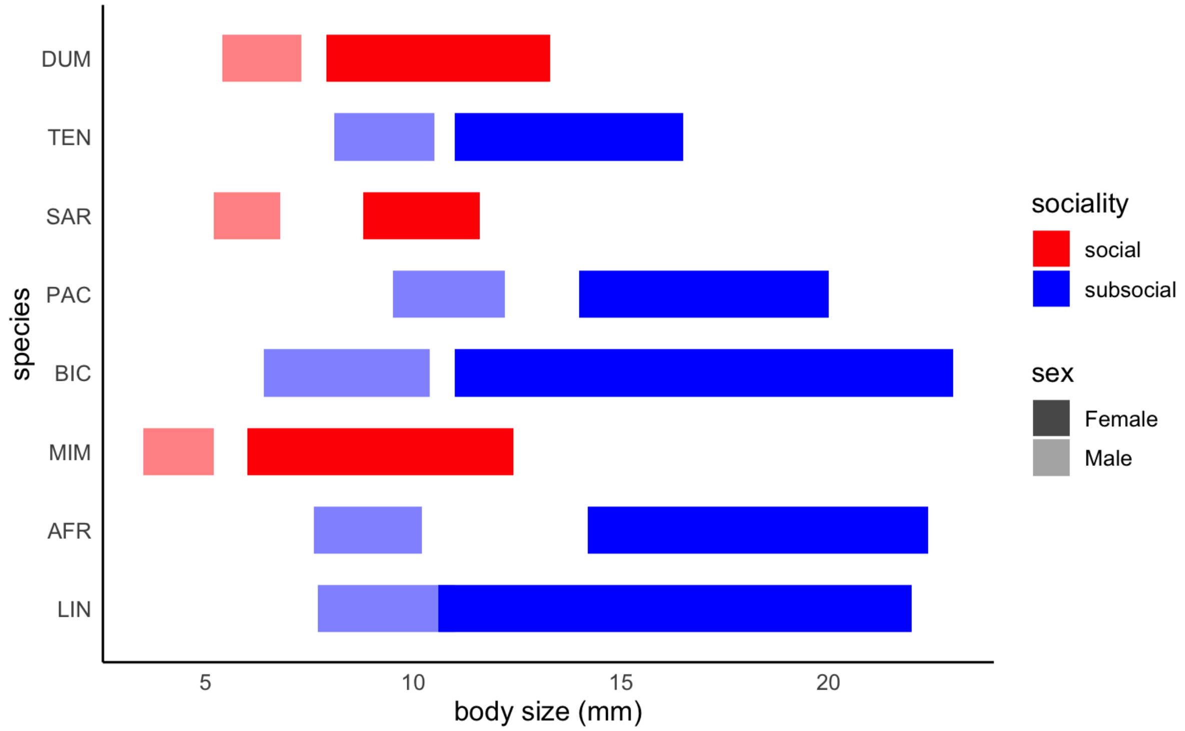

Figure S31. Body-size differences between sexes among Stegodyphus species.

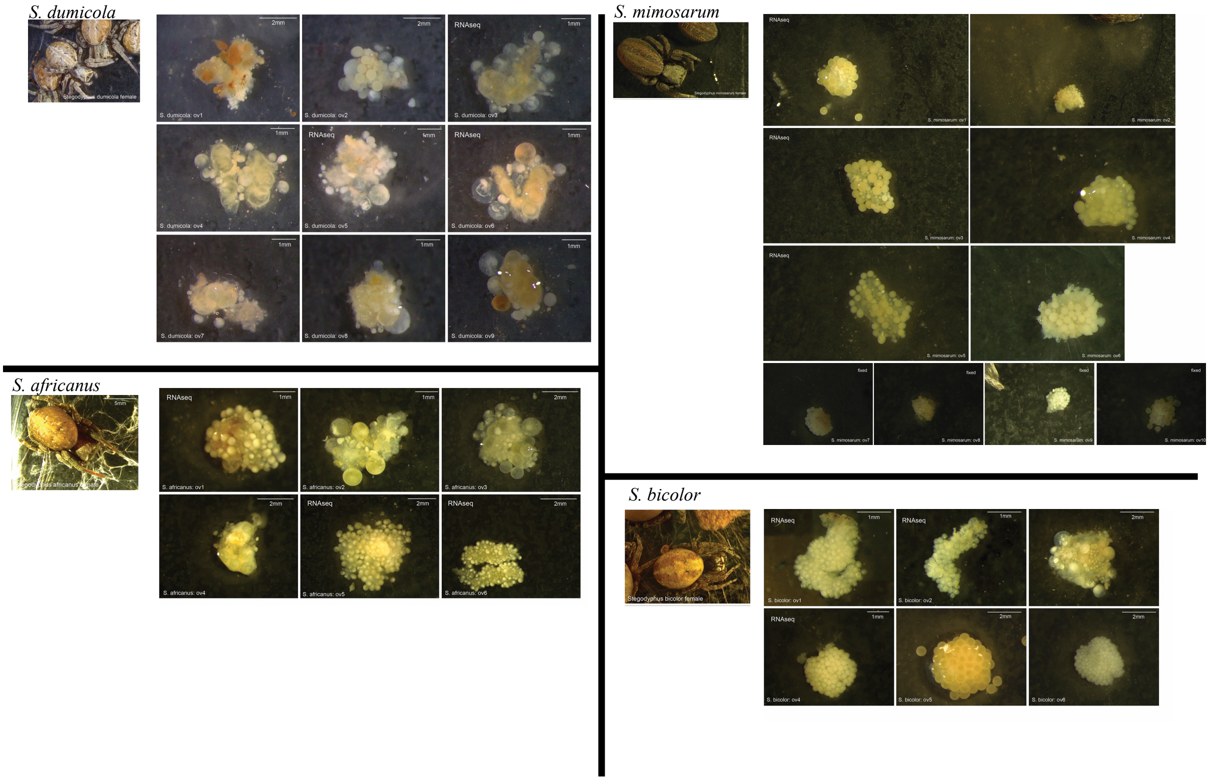

Figure S32. Ovary dissection overview. The selected samples for RNA sequencing is marked with "RNAseq"

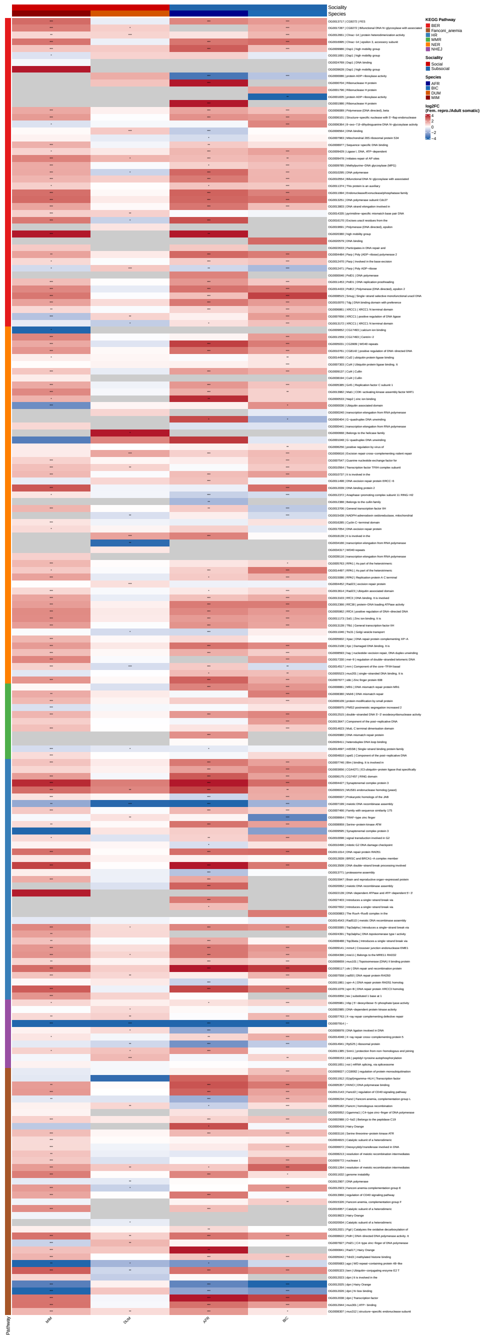

Figure S33: Heatmap of log2 fold change in RNA expression of orthogroups between reproductive (ovary) and somatic tissue for four *Stegodyphus* species, two social (red: *S. mimosarum*, orange: *S. dumicola*) and two subsocial (dark blue: *S. africanus*, blue: *S. bicolor*). Each column represents a differential expression analysis within the given species. The orthogroups are ordered in six KEGG pathways with relation to repair mechanisms, indicated by the annotation color on the left. In the heatmap, red indicates higher expression in the ovary, while blue indicates higher expression in somatic tissue, grey represents an orthogroup not present in a given species genome. significance of the differential expression tests is indicated by asterixes. Orthogroup name is given for each row, followed by common gene name from *drosophila*, if such one exists, as well as a short description of the orthogroup type or function. Expressions were summed across each orthogroup within each species, and differential expression analysis was run on those orthogroups (see methods).

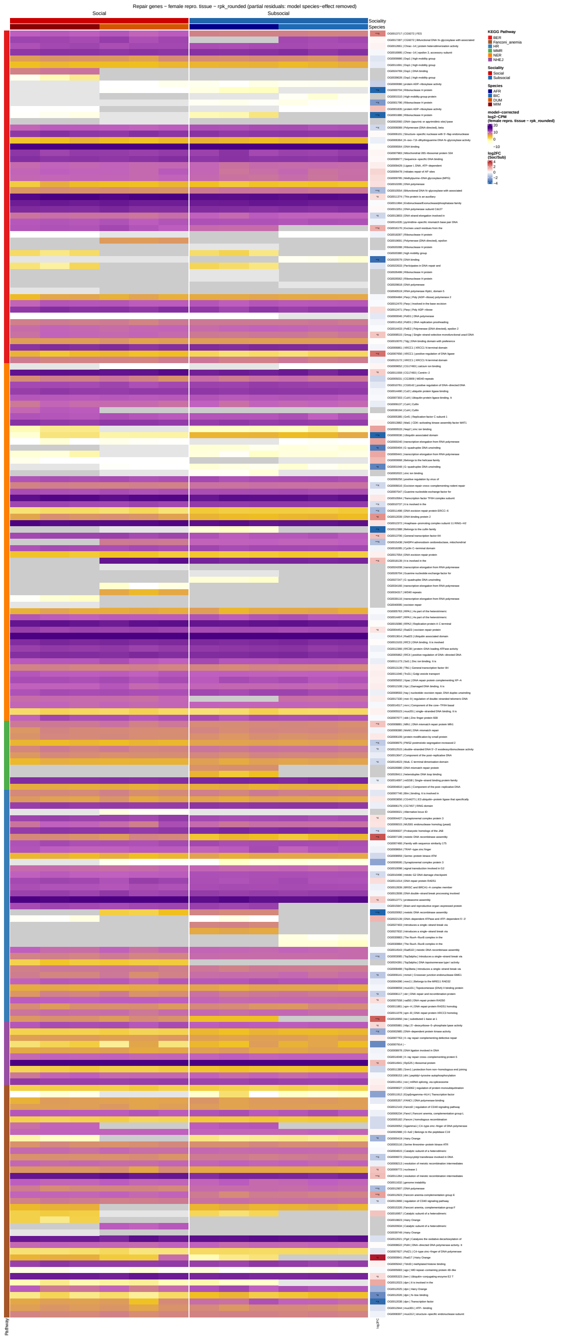

Figure S34: Heatmap of orthogroup RNA expression in reproductive (Ovary) tissue for four *Stegodyphus* species, two social species (red: *S. mimosarum*, orange: *S. dumicola*) and two subsocial species (dark blue: *S. africanus*, blue: *S. bicolor*). The values shown are partial residuals from the differential expression model (log2 CPM), showing what the model sees as potential social-subsocial differences. Dark grey cells indicate orthogroups not annotated in a given species, while light grey indicates orthogroup presence, but no recorded expression. The rightmost column shows log2 fold change for the contrast between social species vs. subsocial species, with red indicating higher expression in social species and blue indicating higher expression in subsocial species and grey indicating insufficient data for testing. The orthogroups are ordered according to KEGG repair pathway (leftmost color annotation), and each orthogroup is annotated with orthogroup name, common *drosophila* name (if it exists) and a short description of the gene orthogroup.

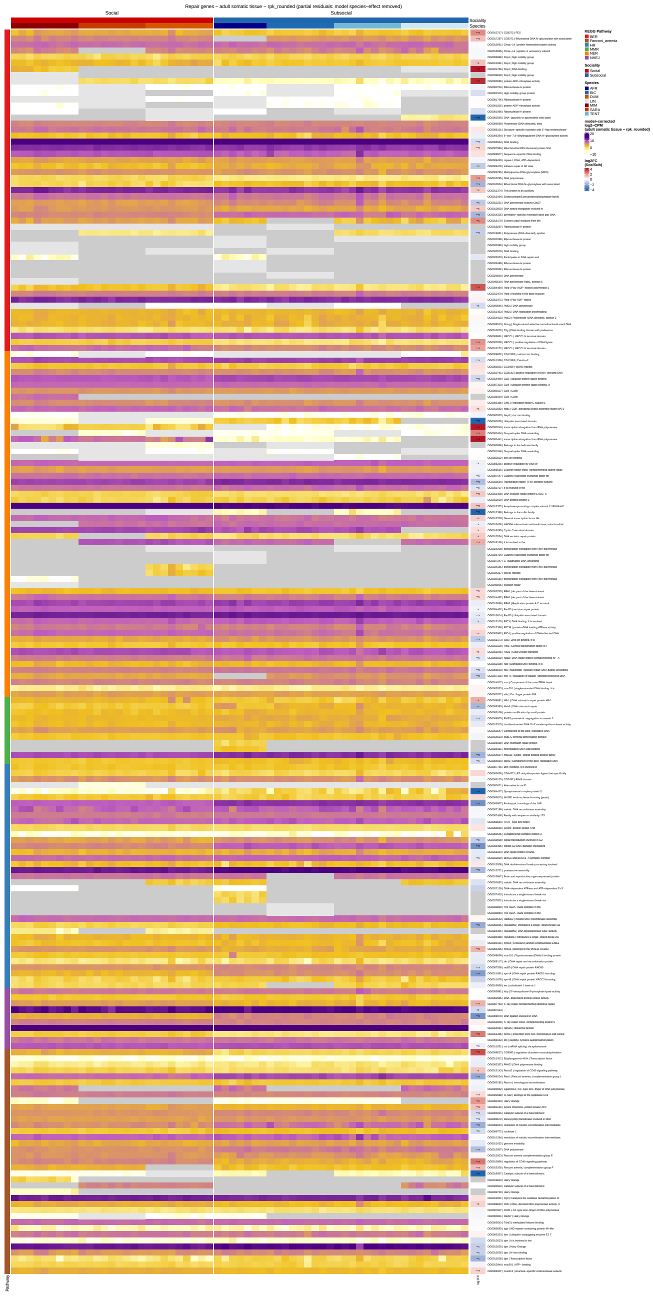

Figure S35: Heatmap of orthogroup RNA expression in adult somatic tissue for seven *Stegodyphus* species, three social species (dark red: *S. mimosarum*, red: *S. sarasinorum*, orange: *S. dumicola*) and four subsocial species (dark blue: *S. africanus*, blue: *S. bicolor*, light blue: *S. tentoriicola*, pale blue: *S. lineatus*). The values shown are partial residuals from the differential expression model (log2 CPM), showing what the model sees as potential social-subsocial differences. Dark grey cells indicate orthogroups not annotated in a given species, while light grey indicates orthogroup presence, but no recorded expression. The rightmost column shows log2 fold change for the contrast between social species vs. subsocial species, with red indicating higher expression in social species and blue indicating higher expression in subsocial species and grey indicating insufficient data for testing. The orthogroups are ordered according to KEGG repair pathway (leftmost color annotation), and each orthogroup is annotated with orthogroup name, common *drosophila* name (if it exists) and a short description of the gene orthogroup.
